# Divergent Cl^−^ and H^+^ pathways underlie transport coupling and gating in CLC exchangers and channels

**DOI:** 10.1101/753954

**Authors:** Lilia Leisle, Yanyan Xu, Eva Fortea, Jason Galpin, Malvin Vien, Christopher A. Ahern, Alessio Accardi, Simon Bernèche

**Affiliations:** Department of Anesthesiology, Weill Cornell Medical College, New York, NY, USA; SIB Swiss Institute of Bioinformatics, University of Basel 50/70 Klingelbergstrasse, 4056 Basel, Switzerland; Biozentrum, University of Basel 50/70 Klingelbergstrasse, 4056 Basel, Switzerland; Department of Physiology and Biophysics, Weill Cornell Medical College, New York, NY, USA; Department of Molecular Physiology and Biophysics, University of Iowa, Iowa City, IA, USA; Department of Biochemistry, Weill Cornell Medical College, New York, NY, USA

## Abstract

The CLC family of anion transporting proteins is comprised of secondary active H^+^-coupled exchangers and of Cl^−^ channels. Both functional subtypes play key roles in human physiology, and mutations causing their dysfunction lead to numerous genetic disorders. Current models suggest that the CLC exchangers do not utilize a classical ‘ping-pong’ mechanism of antiport, where the transporter sequentially interacts with one substrate at a time. Rather, in the CLC exchangers both substrates bind and translocate simultaneously while moving through partially congruent pathways. How ions of opposite electrical charge bypass each other while moving in opposite directions through a shared permeation pathway remains unknown. Here, we use MD simulations in combination with biochemical and electrophysiological measurements to identify a pair of highly conserved phenylalanine residues that form an aromatic pathway, separate from the Cl^−^ pore, whose dynamic rearrangements enable H^+^ movement. Mutations of these aromatic residues impair H^+^ transport and voltage-dependent gating in the CLC exchangers. Remarkably, the role of the aromatic pathway is evolutionarily conserved in CLC channels. Using atomic-scale mutagenesis we show that the electrostatic properties and conformational flexibility of these aromatic residues are essential determinants of channel gating. Our results suggest that Cl^−^ and H^+^ move through physically distinct and evolutionarily conserved routes through the CLC channels and transporters. We propose a unifying mechanism that describes the gating mechanism of CLC exchangers and channels.

## Introduction

The CLC (<u>C</u>h<u>L</u>oride <u>C</u>hannel) family is comprised of Cl^−^ channels and H^+^-coupled exchangers whose primary physiological task is to mediate anion transport across biological membranes (Alessio Accardi, 2015; Thomas J. Jentsch & Pusch, 2018). The human genome encodes for nine CLC homologues, four (CLC-1, -2, -Ka, -Kb) are Cl^−^ channels that reside in the plasma membrane and five (CLC-3 through -7) are 2 Cl^−^:1 H^+^ antiporters that localize to intracellular compartments along the endo-lysosomal pathway. Mutations in five of the human CLC genes lead to genetically inherited disorders of muscle (Thomsen and Becker type *Myotonia congenita*), kidney (Bartter Syndrome Type III and IV, Dent’s disease), bone (Osteopetrosis) and central nervous system (neuronal ceroid lipofuscinosis, retinal degeneration, deafness), highlighting the fundamental roles of CLC channels and transporters in human physiology (T. J. Jentsch, 2008; Stauber, Weinert, & Jentsch, 2012). Several disease-causing mutations occurring in CLC channels and transporters impair the response to the physiological stimuli regulating their activity, such as voltage, pH and nucleotide concentration (Alessio Accardi, 2015; Alekov, 2015; Bignon et al., 2018; Thomas J. Jentsch & Pusch, 2018).

High-resolution structural information on the CLC-ec1 and cmCLC exchangers (Dutzler, Campbell, Cadene, Chait, & MacKinnon, 2002; Feng, Campbell, Hsiung, & MacKinnon, 2010) and CLC-K and CLC-1 channels (E. Park, Campbell, & MacKinnon, 2017; Eunyong Park & MacKinnon, 2018);Wang, 2019 #3114} revealed that both CLC subtypes share a common dimeric architecture, where each monomer forms physically distinct and functionally independent ion permeation pathways. This pathway is defined by 3 anionic binding sites (A. Accardi & Picollo, 2010; Dutzler et al., 2002; Dutzler, Campbell, & MacKinnon, 2003; Feng et al., 2010; E. Park et al., 2017) (Fig. 1A). The external and central sites, S_ext_ and S_cen_, are alternatively occupied by the permeant anions or by the negatively charged side chain of a conserved glutamic acid, Glu_ex_ (Fig. 1A-C). The internal site, S_int_, binds anions weakly and likely serves as a recruitment site of intracellular permeating ions (Lobet & Dutzler, 2006; Picollo, Malvezzi, Houtman, & Accardi, 2009). The presence of a fourth binding site, S_ext*_, in direct contact with the extracellular solution has been proposed based on electrostatic calculations and is supported by electrophysiological data (Faraldo-Gomez & Roux, 2004; Lin & Chen, 2000).

**Figure 1.**
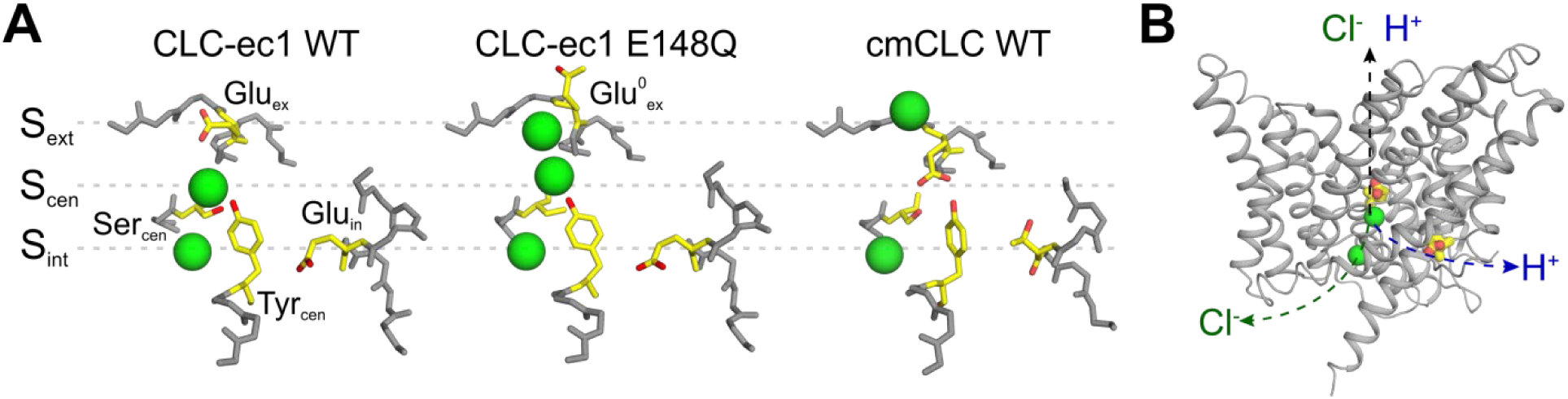
The anion pathway of the CLC Cl^−^/H^+^ exchangers. **(A)** Close up view of the Cl^−^ permeation pathway in three configurations: *left*, Glu_ex_ bound to S_ex_ (CLC-ec1 WT; PDBID: 1OTS); *center*, protonated Glu_ex_ (mimicked by E148Q mutation) reaching out of the ion pathway (PDBID: 1OTU); *right*, Glu_ex_ bound to S_cen_ (cmCLC WT, PDBID: 3ORG). Selected residues are shown as sticks and Cl^−^ ions as green spheres. **(B)** The partially congruent Cl^−^ and H^+^ pathways are shown in CLC-ec1 WT structure (Glu_ex_ and Glu_in_ are shown in yellow, Cl_-_ ions as green spheres).

The CLC Cl^−^ pore can adopt at least three conformations, differentiated by the position and protonation state of Glu_ex_ (Dutzler et al., 2002; Dutzler et al., 2003; Feng et al., 2010) (Fig. 1A). Extensive functional work suggests that cycling through these conformations underlies the Cl^−^/H^+^ exchange cycle in the CLC transporters and gating in the CLC channels (Basilio, Noack, Picollo, & Accardi, 2014; Feng et al., 2010; Feng, Campbell, & MacKinnon, 2012; Khantwal et al., 2016; Lísal & Maduke, 2008; Vien, Basilio, Leisle, & Accardi, 2017). All proposals accounting for the CLC exchange cycle suggest that these transporters do not utilize a classical ‘ping-pong’ mechanism of antiport, where the transporter sequentially interacts with one substrate at a time (Picollo, Xu, Johner, Bernèche, & Accardi, 2012). Rather, the CLCs were proposed to simultaneously bind (Picollo et al., 2012) and translocate Cl^−^ and H^+^ through partially congruent pathways (A. Accardi et al., 2005; Zdebik et al., 2008) (Fig. 1D). The H^+^ pathway is delimited by two glutamic acids that are conserved in the CLC transporters: Glu_in_ serves as the intracellular proton acceptor and is distal from the Cl^−^ permeation pathway, while Glu_ex_ is the extracellular intersection between the H^+^ and Cl^−^ pores (Fig. 1D) (A. Accardi et al., 2005; Zdebik et al., 2008). Several models have been proposed for the exchange cycle of the CLCs (Basilio et al., 2014; Feng et al., 2010; Khantwal et al., 2016; C. Miller & Nguitragool, 2009), however none provided a molecular mechanism describing how the Cl^−^ and H^+^ ions bypass each other while moving in opposite directions through the permeation pathway. These proposals share the critical assumption that protonation of Glu_ex_ within the pathway destabilizes its binding to S_cen_ and/or S_ext_, favoring its exit from the pathway. While this mechanism readily explains outward H^+^ transfer, it predicts that when the transporter is mediating H^+^ influx, a protonated and neutral Glu^0^_ex_ outcompetes the negatively charged Cl^−^ ions bound to the anion-selective S_ext_ and S_cen_ sites. This is an energetically unfavorable transition, which should result in intrinsic rectification of transport. Indeed, free-energy calculations show that a protonated Glu_ex_ encounters a high energy barrier to enter the Cl^−^ permeation pathway (Kuang, Mahankali, & Beck, 2007). In contrast, the CLC exchangers function with comparable efficiency in the forward and reverse directions (De Stefano, Pusch, & Zifarelli, 2013; L. Leisle, Ludwig, Wagner, Jentsch, & Stauber, 2011; Matulef & Maduke, 2005), suggesting that the entry and exit of the protonated Glu_ex_ from the Cl^−^ pathway are equally favorable. Further, the mechanisms regulating the release of ions from S_ext_ and S_cen_ and their coupling to the movement of Glu_ex_ and of H^+^ through the protein are unknown. While biochemical evidence suggests that ion release is rate-limited by a conformational step (Picollo et al., 2009), no release mechanism has been identified. Therefore, two key mechanistic features at the heart of the H^+^:Cl^−^ exchange mechanism of the CLCs, the pathways and coupling mechanism of the substrates, remain unknown.

Here we combined molecular dynamics simulations with biochemical and electrophysiological measurements, and atomic mutagenesis to investigate the mechanism of H^+^/Cl^−^ exchange. We find that, contrary to previously proposed models, a protonated Glu_ex_ does not move through the Cl^−^ pore. Rather, we identify two highly conserved phenylalanine residues that form an aromatic slide which allows the protonated (neutral) Glu_ex_ to move to and from S_cen_ without directly competing with Cl^−^ ions for passage through the anion-selective pathway. Further, we show that the rotational movement of the central phenylalanine residue, that enables the formation of the aromatic slide, regulates ion movement within the pathway, providing the molecular mechanism for the coupled exchange of H^+^ and Cl^−^ by the CLC exchangers. Mutating these residues in prokaryotic and mammalian CLC exchangers severely impairs transport indicating that the role of these aromatic side chains is evolutionarily conserved. Since these phenylalanine residues are highly conserved throughout the CLC family, we hypothesized the role of the aromatic slide might be evolutionary conserved also between CLC channels and exchangers. Indeed, we found that they are critical determinants of gating of the prototypical CLC-0 channel. Using atomic-scale mutagenesis, we probed how the aromatic slide residues interact with Glu_ex_ in CLC-0, and found that the central phenylalanine interacts electrostatically with the gating glutamate, and that its conformational rotation is necessary for channel opening. We propose a novel mechanism for CLC mediated H^+^:Cl^−^ exchange, where the Cl^−^ and H^+^ pathways are distinct and intersect only near the central ion binding site.

## Results

### Molecular dynamic simulations suggest F357 controls entry and release of Cl^−^ from S_cen_

We used molecular dynamics simulations to probe the energetic landscape of ion movement through the permeation pathway of CLC-ec1 to ask whether it is regulated by conformational rearrangements of the pore. The first state we considered is one with Glu_ex_ protonated and out of the pathway, an E148Q-like state (Dutzler et al., 2003) (Fig. 1A), so that all binding sites are accessible to ions. The first Cl^−^ ion to reach the pore preferably binds to S_cen_ or S_ext_, this remains true when another ion is bound to S_ext_* (Fig. 2A, dashed line (i), 2B). When a Cl^−^ occupies S_int_, the second ion can occupy S_cen_ or S_ext_, though binding to S_cen_ is less favorable by 2 kCal mol^−1^ (Fig. 2A, dashed line (ii), 2B). Unexpectedly, our calculations suggest that simultaneous binding of Cl^−^ to S_cen_ and S_ext_ does not correspond to a local free energy minimum, i.e. to a (meta-)stable state. The doubly occupied state, identified by an asterisk on Fig. 2A, is unfavorable by 4 to 6 kCal mol^−1^ in comparison to the binding of a single ion to S_ext_ or to the doubly occupied configuration with ions in S_int_ and S_ext_.

**Figure 2.**
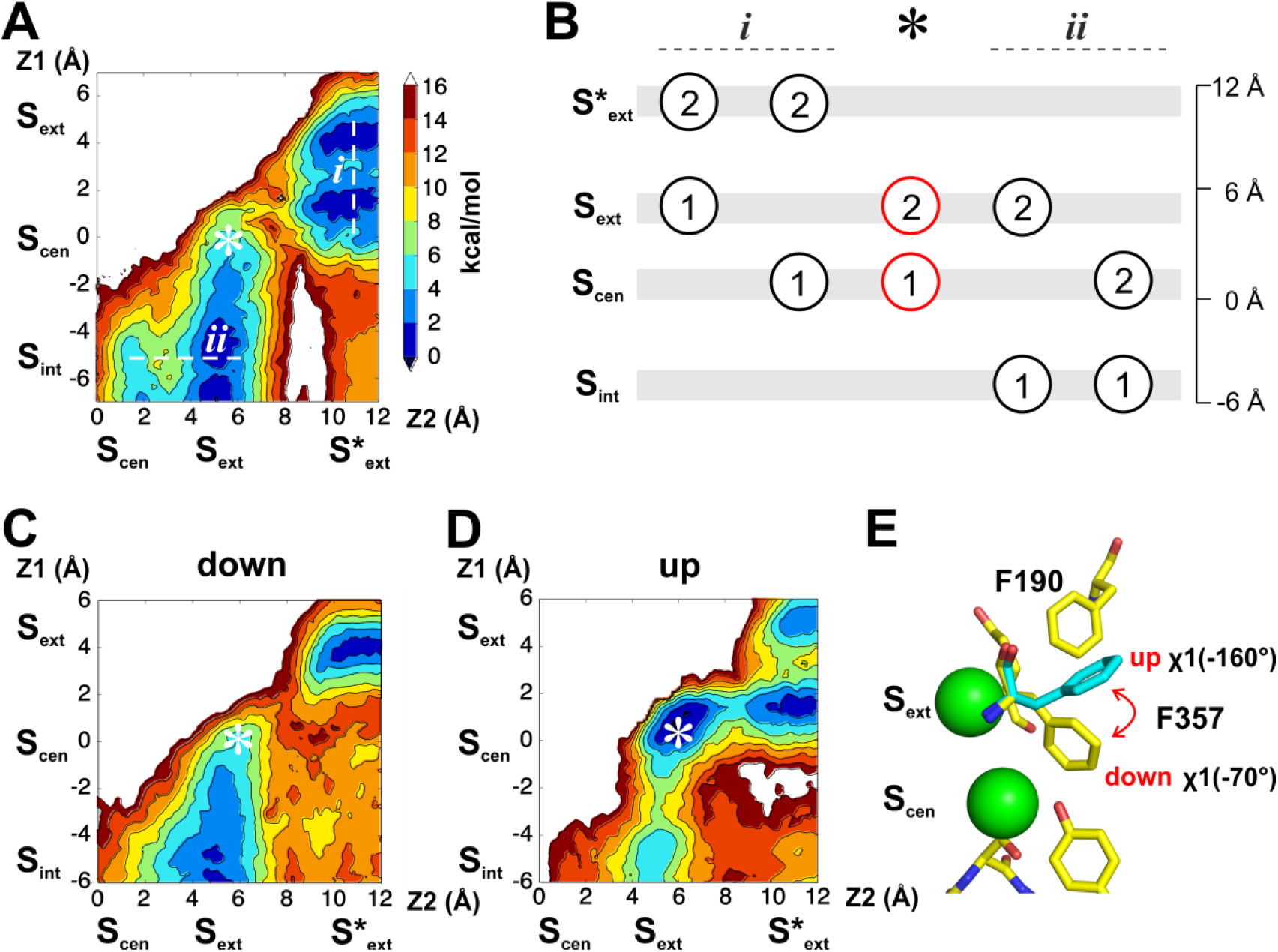
Energetic landscapes of ion movement through the pathway of CLC-ec1 reveal two rotameric states of F357 that correlate with ion occupancy. (**A**) The PMF calculation describes the energetics of two Cl^−^ along the permeation pathway, with the protonated Glu^0^_ex_ positioned on the extracellular side of the pore (E148Q-like conformation). For both ions, the reaction coordinate consists in the position along the Z-axis (Z1, Z2). The position of the different binding sites along the Z-axis is indicated. The Z=0 coordinate is an arbitrary point defined as the center of mass of backbone atoms around S_cen_. The asterisk indicates the position of the doubly occupied state S_cen_/S_ext_. Each color of the iso-contoured map represents a dG of 2 kCal mol^−1^, as indicated by the color scale. (**B**) The scheme illustrates the key occupancy states. The labels at the top refer to the positions indicated in panel A. The doubly occupied state S_cen_/S_ext_ is essential for ion transport but is not observed as a stable state in the PMF calculation of panel A. (**C-D**) The PMF calculation was repeated with a harmonic restraint applied to the side chain of residue F357 to maintain *χ*1 around -70° (down conformer, C) or -160° (up conformer, D). Simultaneous Cl^−^ binding to S_ext_ and S_cen_ becomes energetically favorable under the ‘up’ conformer. (**E**) F357 side chain exists in an equilibrium between two rotameric states (‘up’, *χ*1=-160°; ‘down’, *χ*1=-70°).

Since current models postulate a state with simultaneous occupancy of S_cen_ and S_ext_, we set out to investigate what gives rise to this energetic barrier. Analyzing the conformational sampling underlying the PMF calculations, we noted that simultaneous occupancy of S_cen_ and S_ext_ correlated with fluctuations of the rotameric state of F357 (Fig. 2 – Supplement 1A, B). This residue forms part of the Cl^−^ permeation pathway of the CLC channels and transporters by coordinating ions in S_cen_ and S_ext_ with its backbone amide (E. Park et al., 2017). In our simulations, we find that the F357 side chain exists in equilibrium between two rotameric states with *χ*1 angles of -160° (‘up’), as seen in the crystal structures of WT and mutant CLC-ec1 (Dutzler et al., 2002; Dutzler et al., 2003), and of -70° (‘down’) (Fig. 2E). To test whether this conformational rearrangement affects Cl^−^ permeation, we restrained *χ*1 (F357) to the ‘up’ or ‘down’ rotamers and determined the energetic landscape of ion binding in these conformations (Fig. 2C, D). We find that when F357 is in the ‘down’ state, a free energy barrier of ∼12 kCal mol^−1^ opposes ion movement within the pore, and the S_cen_/S_ext_ double occupancy state remains unstable (Fig. 2C). In contrast, when F357 is constrained to the ‘up’ conformer, two Cl^−^ ions can simultaneously bind to S_cen_ and S_ext_, and the different stable states along the permeation pathways are separated by free energy barriers of ∼2-4 kCal mol^−1^ (Fig. 2D). Reciprocally, the ion occupancy state directly impacts the conformation of F357: the ‘down’ state is favored when no ions occupy the pathway or when only S_ext_ is occupied (Fig. 2 – Supplement 1C, D), ion occupancy of S_cen_ or of S_cen_ and S_ext_ simultaneously favors the ‘up’ state of F357 (Fig. 2 – Supplement 1E, F). This suggests that the F357 transition between the ‘up’ and ‘down’ rotamers is critical for allowing ion permeation. To test this hypothesis, we asked whether the dynamics of F357 are affected by the introduction of a crosslink between A399 and A432 (Fig. 2 – Supplement 2A), which inhibits transport of CLC-ec1 by preventing a conformational rearrangement involved in coupling between the intra- and extra-cellular gates (Basilio et al., 2014). We found that constraining the relative movements of A399 and A432 inhibits the transition of F357 between its rotamers by increasing the transition free energy barrier by about 6 kCal mol^−1^ (Fig. 2 – Supplement 2), suggesting that this transition is part of the transport cycle. Taken together, these results suggest that the rearrangement of F357 is a critical determinant of the energy barrier height for ion movement within the CLC pore: the ‘up’ conformer of F357 is compatible with ion transport while its ‘down’ conformer disfavor ion transitions.

### Rotation of F357 enables the formation of an aromatic slide

We next asked how Cl^−^ and Glu_ex_ interact along the transport cycle. As in the previous section, we first considered states in which Glu_ex_ is protonated (Glu^0^_ex_) and outside the Cl^−^ pathway in an E148Q-like conformation (Fig. 3A). Analysis of the conformational sampling of the PMFs presented in Fig. 2 reveals that, in this configuration, Glu^0^_ex_ is stabilized by the carboxylate group of D54 via a water molecule (Fig. 3A) or by a hydrogen bond with the backbone of A189 (Fig. 3B). The simulations also reveal that the carboxylate group of Glu^0^_ex_ rarely visits S_ext_ and rather interacts with the aromatic ring of F190, even if S_ext_ is free of Cl^−^ (Fig. 3C).

**Figure 3.**
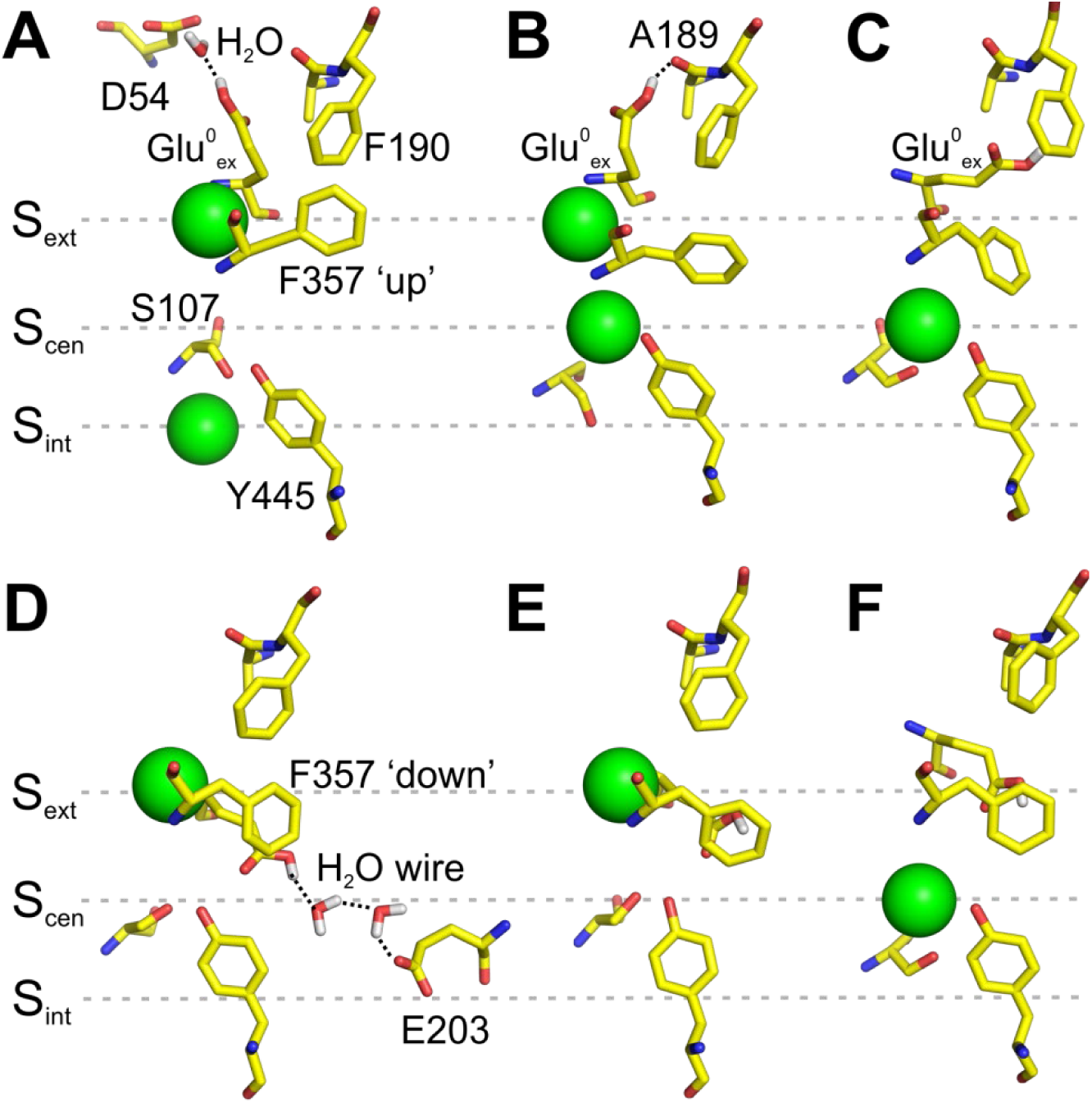
Interaction of Glu_ex_ with F190 and F357. (**A-C**) Conformations of the pore extracted from the PMF calculation presented in Figure 2A, in which Glu^0^_ex_ is initially placed on the extracellular side of the pore. Glu^0^_ex_ is part of a hydrogen bond network involving D54 (**A**) or A189 (**B**). Glu^0^_ex_ can also interact with the side chain of F190 (**C**). (**D-F**) Conformations of the pore extracted from a 1D PMF describing the binding of a Cl^−^ to the pore when Glu^0^_ex_ is initially bound to S_cen_ (see Figure 3 – Supplement 1). A proton wire is spontaneously formed between Glu_ex_ (E148) and Glu_in_ (E203), which are bridged by two water molecules (**D**). Glu^0^_ex_ can form a dipole-*π* interaction with F357 aromatic side chain, leaving S_cen_ empty (**E**). Cl^−^ moves from S_ext_ to S_cen_, while the side chain of Glu^0^_ex_ is stabilized outside the ion pathway by its interaction with F357, and in proximity of F190 (**F**).

We then considered states in which Glu_ex_ occupies S_cen_. We calculated the PMF describing the binding of Cl^−^ to S_ext_ for both the charged and protonated forms of Glu_ex_ (Fig. 3 – Supplement 1). The PMFs show that the binding of Cl^−^ to S_ext_ requires the protonation of Glu_ex_, in agreement with our previous work suggesting synergistic binding of Cl^−^ and H^+^ (Picollo et al., 2012). Interestingly, in the case of the protonated Glu_ex_, a free energy well is also observed at the level of S_cen_, initially occupied by the carboxylate group of Glu^0^_ex_. Inspection of the sampled structures reveals that, when S_ext_ is occupied by a Cl^−^, the Glu^0^_ex_ only partially occupies S_cen_. Its carboxylate group moves sideway toward F357 and away from the Cl^−^ pathway. Two key conformations are observed. A first one in which a proton wire composed of two water molecules forms spontaneously between Glu^0^_ex_ (E148) and Glu_in_ (E203) (Fig. 3D), potentially allowing the deprotonation of Glu^0^_ex_. In this conformation, the carboxylate group of Glu^0^_ex_ also forms a hydrogen bond with Y445. A second conformation reveals the possibility for Glu^0^_ex_ to form a π-dipole interaction with the aromatic ring of F357 (Fig. 3E). The displacement of Glu^0^_ex_ toward F357 allows the bound Cl^−^ to reach S_cen_ (Fig. 3F). The upward movement of F357 brings the carboxylate group of Glu^0^_ex_ in the vicinity of F190. These calculations suggest that the conformational rearrangement of F357 enables the formation of an aromatic slide through which a protonated Glu^0^_ex_ can move to and from S_cen_ without having to compete with the bound ions in the ion pathway.

### The aromatic slide residues are essential for Cl^−^:H^+^ coupling and exchange in CLC-ec1

Our molecular dynamics simulations suggest that F190 and F357 play a critical role in determining Cl^−^/H^+^ coupling and control a rate-limiting barrier for ion transport in CLC-ec1. To test these hypotheses, we mutated them to alanine and determined the unitary transport rate and stoichiometry of the Cl^−^/H^+^ exchange cycle. Both mutations slow the turnover rate and degrade the exchange stoichiometry (Fig. 4). The F190A mutant slows transport ∼2-fold (Fig. 4B, D Fig.4 – Suppl. 1C), while it severely impairs the transport stoichiometry to ∼8.6:1 (Fig. 4A, E, Fig.4 – Suppl. 1C). The F357A mutant drastically reduces the transport rate ∼5.4-fold and alters the Cl^−^ :H^+^ stoichiometry to ∼4.3:1 (Fig. 4C-E, Fig.4 – Suppl. 1C). As the F357 backbone lines S_cen_ and S_ext_ (Dutzler et al., 2002) we tested whether an alanine substitution affects the integrity of the binding sites in two ways. First, we used isothermal titration calorimetry (ITC) to measure Cl^−^ binding to the F357A mutant, and found that it has a WT-like affinity of ∼0.7 mM (Fig. 4 – Supplement 1A). Second, we introduced the F357L substitution, which preserves the hydrophobicity and volume of side chain but removes the aromatic ring, and found that it affects the turnover rate and exchange stoichiometry like the F357A (Fig. 4 – Suppl. 1B, C). Thus, the effects of the F357A mutant reflect the absence of the aromatic side chain rather than a structural disruption of the ion pathway. Taken together our functional results support the insights from our MD simulations that F190 and F357 play a key role in coupling and transport of CLC-ec1. Remarkably, F190 and F357 are among the highest conserved residues throughout the CLC family, respectively at ∼94% and ∼76% (Fig. 4 – Suppl. 2), suggesting that their functional role might be evolutionarily conserved. In the remainder of this work we will refer to these residues across different CLC homologues as Phe_ex_ (F190 in CLC-ec1) and Phe_cen_ (F357 in CLC-ec1).

**Figure 4.**
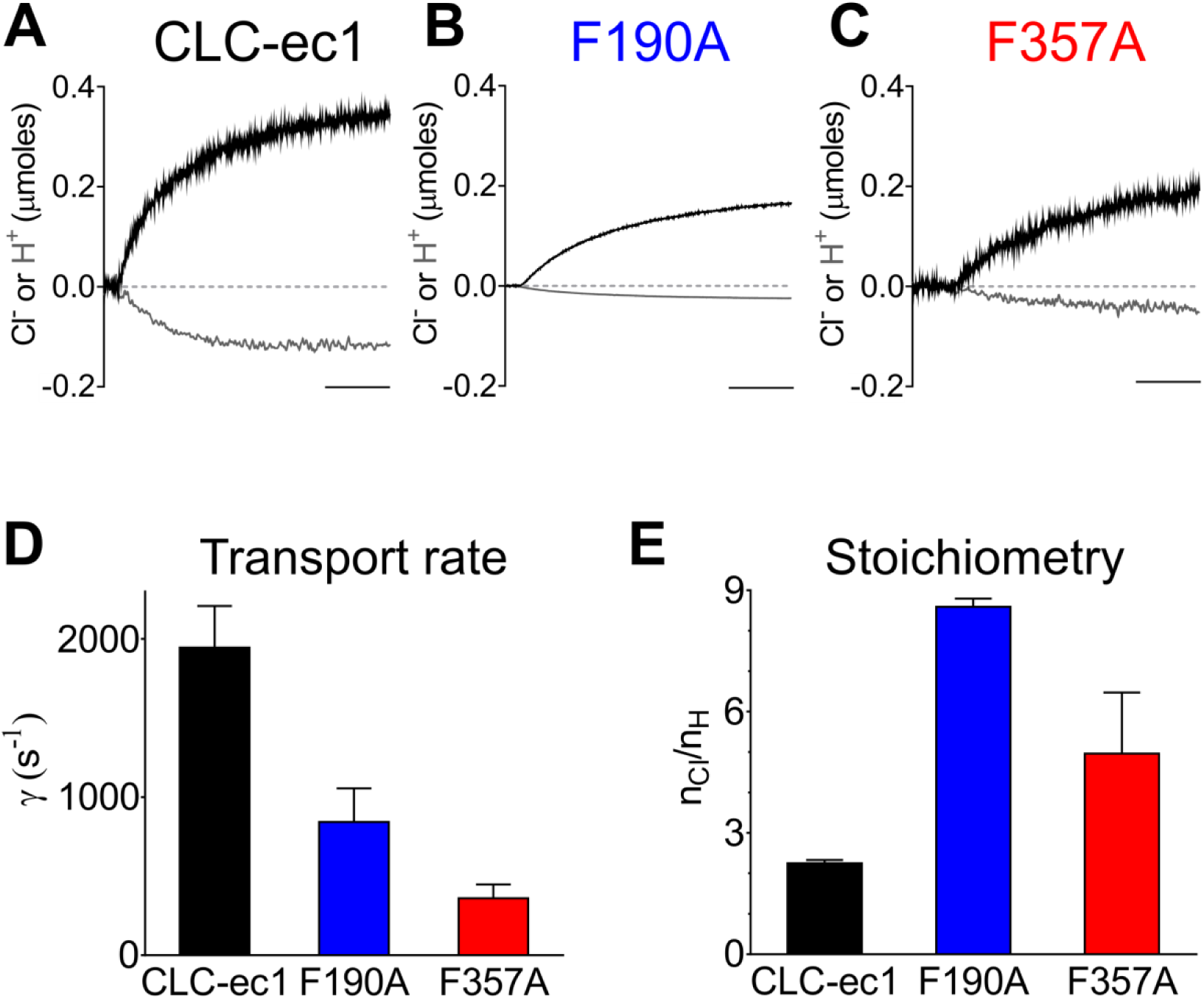
Phe_cen_ (F357) and Phe_ex_ (F190) determine the absolute transport rate and Cl^−^/H^+^ coupling stoichiometry in CLC-ec1. **(A-C)** Representative time course of Cl^−^ (black) and H^+^ (gray) transport recordings of purified CLC-ec1 WT (A), F190A (B) and F357A (C) reconstituted into liposomes. *Scale bar*, 25 s. **(D-E)** Average turnover rate (**D**) and transport stoichiometry (**E**) of CLC-ec1 WT (black), F190A (blue) and F357A (red).

### The role of Phe_ex_ and Phe_cen_ is conserved in mammalian transporters

We asked whether the role of Phe_ex_ and Phe_cen_ is conserved in the mammalian CLC exchangers. The mammalian CLC-5 and CLC-7 exchangers are activated by depolarizing voltages (Fig. 5A, Fig. 5 - Suppl. 2A), with their voltage-dependence arising from the movement of the permeating ions and of Glu_ex_ through the electric field of the CLC transport pathway (Smith & Lippiat, 2010; Zifarelli, De Stefano, Zanardi, & Pusch, 2012). The G-V relationship of CLC-7 is well described by a Boltzmann function with a V_0.5_ of ∼+90 mV, and the currents display slow, voltage-dependent gating relaxation kinetics (Fig. 5)(L. Leisle et al., 2011). We asked how alanine substitution of Phe_ex_ (F301A) and Phe_cen_ (F514A) would affect gating of CLC-7 (Fig. 5). The average steady state currents of F301A and F514A at +90 mV are reduced by ∼25% and ∼50%, respectively, compared to WT (Fig. 5D), in line with the reduced transport rates observed for CLC-ec1 (Fig. 4D). The F301A mutant is nearly voltage independent between -80 and +90 mV (Fig. 5B, E) and so are its gating kinetics (Fig. 5 – Suppl. 1). Conversely, the F514A mutant results in voltage-dependent currents, but with a left-shifted G-V relationship by ∼50 mV (Fig. 5C, E) and with accelerated gating kinetics (Fig. 5 – Suppl. 1). The corresponding mutations in the ClC-5 exchanger, F255A (Phe_ex_) and F455A (Phe_cen_), have qualitatively similar effects to those seen in CLC-7: the F255A mutant drastically reduces voltage-dependent gating, resulting in measurable currents at negative voltages, and the activation threshold of the F455A mutant is left-shifted (Fig. 5 - Suppl. 2). However, the fast gating kinetics and extreme rectification of this homologue prevent a reliable determination of the G-V relationship and thus a quantification of the effects. Taken together, our results show that Phe_ex_ and Phe_cen_ play a key role to enable the movement of Cl^−^ ions and Glu_ex_ in and out the anion pathway. We hypothesize that these residues interact with the permeating ions and Glu_ex_ via direct and electrostatic interactions.

**Figure 5.**
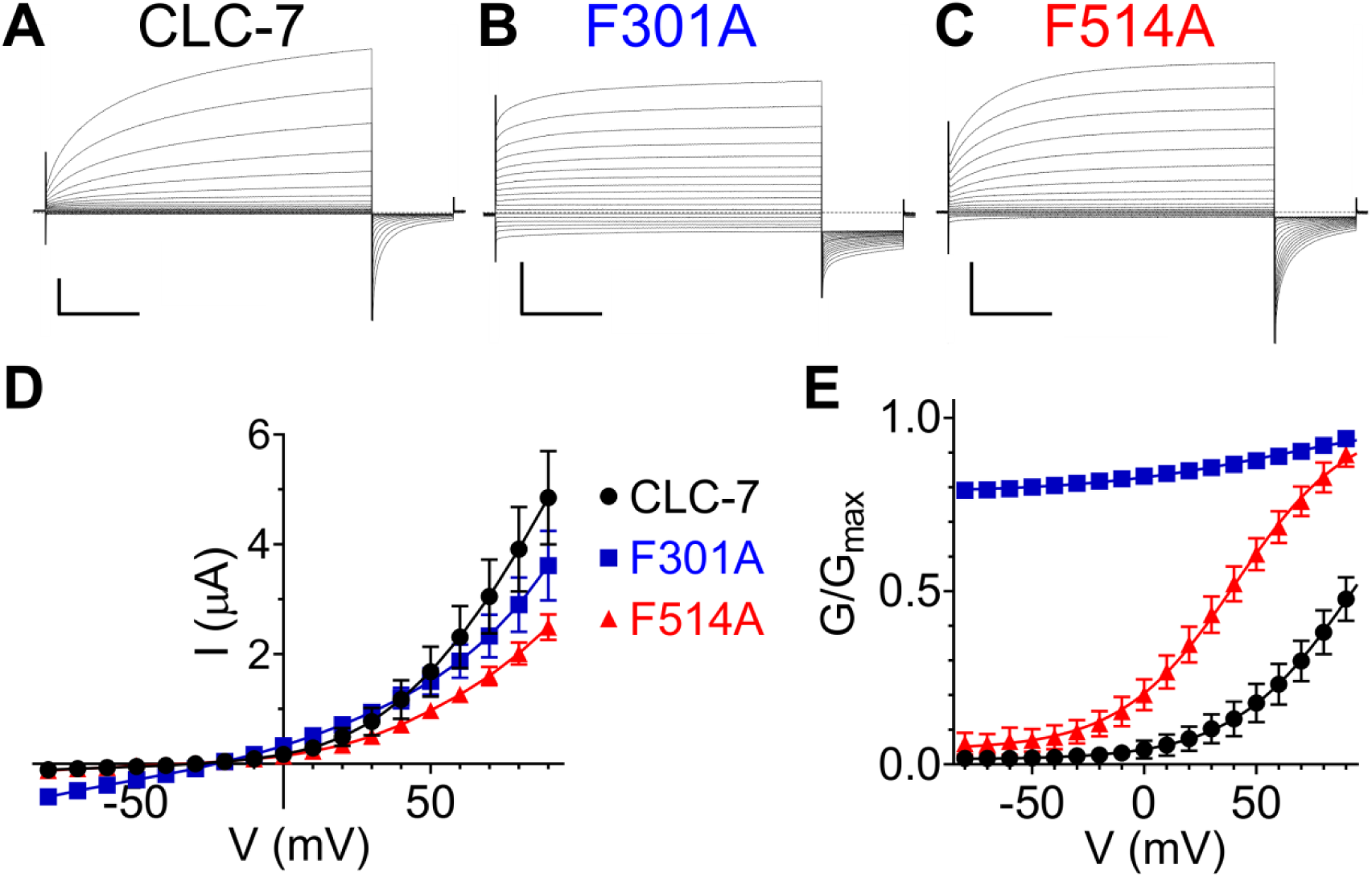
Role of Phe_cen_ (F514) and Phe_ex_ (F301) in the CLC-7 exchanger. **(A-C)** Representative TEVC current recordings of CLC-7 WT (A), F301A (B) and F514A (C). For voltage clamp protocol see *Methods* section. *Horizontal scale bar*, 0.5 s; *vertical scale bar*, 1 μA. **(D)** Steady state I-V relationships for CLC-7 WT (black), F301A (blue) and F514A (red). Symbols represent the average of (N(WT)=7; N(F301A)=18; N(F514A)=12) independent experiments. Solid line holds no theoretical meaning. The mean current amplitudes at +90 mV are I(WT)=4.9±0.9 μA; I(F301A)=3.2±0.5 μA and I(F514)=2.5±0.2 μA. **(E)** G-V relationships for WT and mutant CLC-7 determined from the initial values of tail currents (see *Methods*). Symbols represent the average of 7-18 independent experiments (as in (D)). Solid line is a fit to a Boltzmann function with an offset. Values are reported as mean ± S.E.M, error bars are not shown where they are smaller than the symbol size. Values for the fit parameters and number of repeats are reported in Supplementary Table 1.

### Phe_ex_ and Phe_cen_ are key determinants of CLC channel gating

To probe whether the aromatic slide also plays a similar role in determining the voltage dependent gating of the CLC channels, we mutated Phe_ex_ (F214) or Phe_cen_ (F418) to alanine in the prototypical CLC-0 channel. Opening of CLC-0 channels is regulated by two processes: individual pores open and close independently in a process called single-pore gating and cooperatively during common-pore gating (Alessio Accardi, 2015; Thomas J. Jentsch & Pusch, 2018). Single-pore gating is thought to entail rearrangements of Glu_ex_ similar to those underlying the exchange cycle of the transporters, while the mechanistic underpinnings of common-pore gating remain poorly understood (Alessio Accardi, 2015; Thomas J. Jentsch & Pusch, 2018). In the CLC-0 channel, single-pore gating is activated by depolarizing voltages (Fig. 6A, D) while common-pore gating is hyperpolarization activated and occurs on much slower time scales (Fig. 6E, H). In the F214A and F418A mutants, the single-pore gate is nearly constitutively open, with nearly voltage-independent gating relaxations and G-V relationships (Fig. 6B-D). The voltage dependence of the common-pore gate is also affected in these mutants, as the minimal open probability increases, the gating charge decreases and V_0.5_ shift to more positive values (Fig. 6E-H, Supplementary Table 1). These results suggest that Phe_ex_ and Phe_cen_ are shared determinants of the voltage dependence of the common- and single-pore gating processes of CLC-0. This is consistent with our MD simulations on the CLC-ec1 exchanger, where the carboxyl group of Glu^0^_ex_ interacts with the aromatic-slide residues electrostatically (Fig. 3). However, interpretation of the effects of alanine substitutions, or by any substitution with canonical amino acids, is difficult as these replacements simultaneously affect multiple side-chain properties such as volume, hydrophobicity and electrostatics. To circumvent these limitations, we used non-canonical amino acid (ncAA) mutagenesis to selectively manipulate specific properties of the phenylalanines comprising the aromatic slide (Fig. 7) using three derivatives: Cyclohexylalanine (Cha), 2,6diFluoro-Phenylalanine (2,6F_2_-Phe) and 2,6diMethyl-Phenylalanine (2,6diMeth-Phe; Fig. 7A). In Cha, the benzene ring is replaced by the non-aromatic cyclohexane ring, rendering the side chain electroneutral, while minimally altering size and hydrophobicity (Ahern, Eastwood, Lester, Dougherty, & Horn, 2006; Mecozzi, West, & Dougherty, 1996) (Fig. 7A). In 2,6F_2_-Phe, the fluorine atoms at positions C2 and C6 withdraw π electrons from the face of the benzene ring which makes the edges close to the backbone electronegative and the distal edges electropositive, and, moreover, neutralizes the negative face of the aromatic ring (Ahern et al., 2006; Mecozzi et al., 1996) (Fig. 7A). Importantly, the hydrophobicity of the residue is not affected as benzene and hexa-fluoro-benzene have similar logP values (Leo, Hansch, & Elkins, 1971). Finally, the methyl groups at positions C2 and C6 in 2,6diMeth-Phe (Fig. 7A) restrict the rotameric conversion of the aromatic side chain around the χ1 angle (Harrison, Pasternak, & Verdine, 2003; Li et al., 2007).

**Figure 6.**
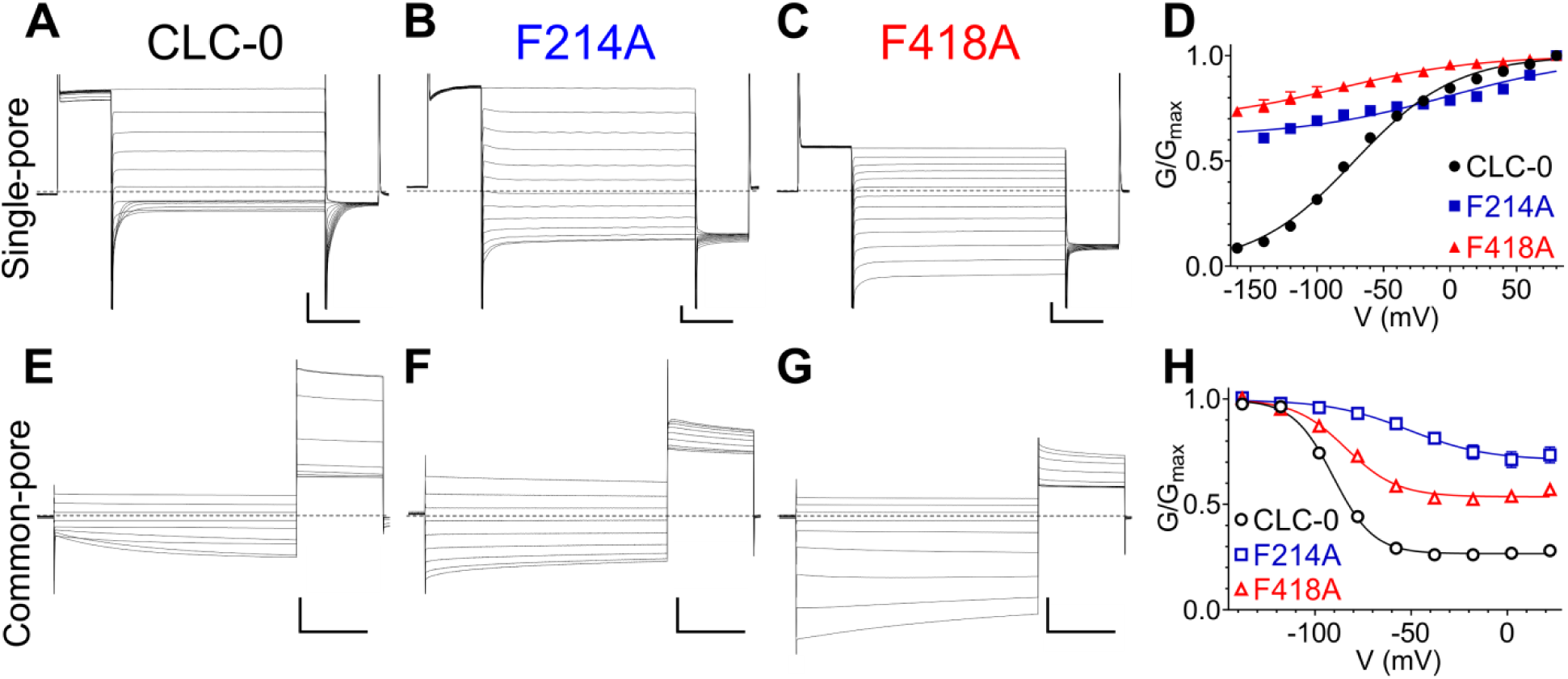
Phe_cen_ (F418) and Phe_ex_ (F214) contribute to voltage dependence of single- and common-pore gate of the CLC-0 channel. **(A-C, E-G)** Representative TEVC current recordings of CLC-0 WT (**A, E**), F214A (**B, F**) and F418A (**C, G**) evoked by single-pore (**A-C**) or common-pore (**E-G**) gating protocols (see *Methods*). *Horizontal scale bar*, 50 ms (single-pore gate) or 2 s (common-pore gate); *vertical scale bar*, 4 μA for CLC-0 WT and 2 μA for F214A and F418A. **(D, H)** Normalized G-V relationships of the single (**D**, filled symbols) and common pore (**H**, empty symbols) gating processes of CLC-0 WT (black circles), F214A (blue squares) and F418A (red triangles). Solid lines represent fits to a Boltzmann function (Eq. 1). Values are reported as mean ± S.E.M, error bars are not shown where they are smaller than the symbol size. Values for the WT fit parameters and number of repeats for all conditions are reported in Supplementary Table 1.

**Figure 7.**
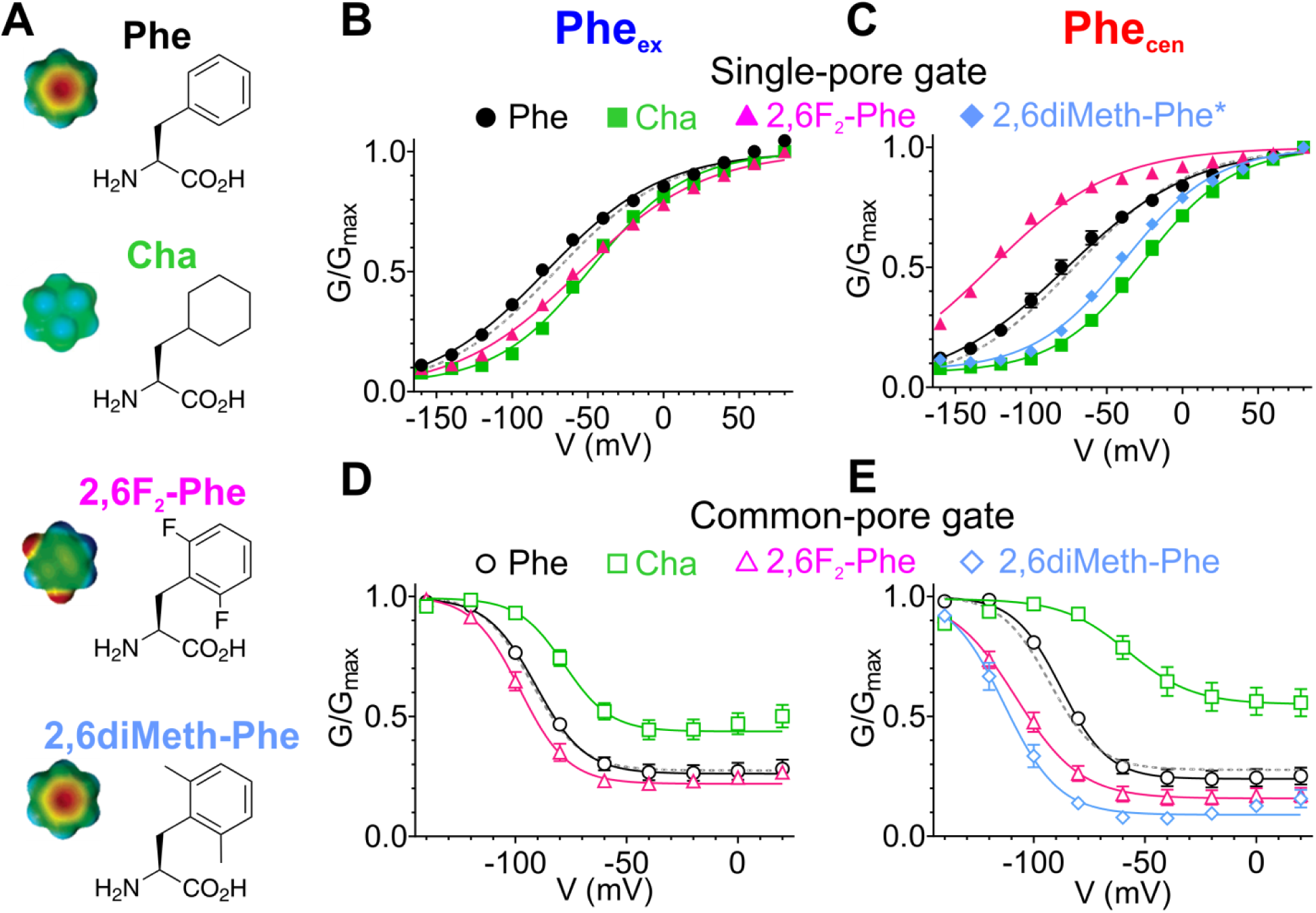
Atomic scale mutagenesis confirms importance of π-electron distribution and rotational movement of Phe_cen_ for CLC-0 gating processes. (**A**) Phenylalanine (Phe) and non-canonical Phe derivatives used in this study: *Cha*, Cyclohexylalanine; *2,6F_2_-Phe*, 2,6Fluoro-Phenylalanine; *2,6diMeth-Phe*, 2,6diMethyl-Phenylalanine. Right panel: stick representation of the amino acids, left panel: surface electrostatic potential of benzene and its derivatives with red and blue corresponding to -20 and +20 kCal/mol, respectively (Mecozzi et al., 1996). The surface electrostatic potential of 2,6diMeth-Phe is assumed similar to Phe because methyl group substitutions do not withdraw electrons from the benzene ring. (**B, C**) Normalized G-V relationships of the single (**B, C**) and common (**D, E**) pore gating processes for CLC-0 with the following replacements at Phe_ex_ (**B, D)** and Phe_cen_ (**C, E**): Phe (black circles), Cha (green squares), 2,6F_2_-Phe (pink triangles) and 2,6diMeth-Phe (cyan diamonds). WT G-V curves (from Fig. 6) are shown as gray dashed lines for reference. Solid lines represent fits to a Boltzmann function with an offset (see *Methods*, Equation 1). Note that the G-V data for the F418X+2,6diMeth-Phe was obtained on the background of C212S mutant (2,6diMeth-Phe*) to isolate its effects on the single-pore gating process. The effects of F418X+2,6diMeth-Phe substitution on the WT background are shown in Fig. 7 - Suppl. 2. Values are reported as mean ± S.E.M, error bars are not shown where they are smaller than the symbol size. Values for the fit parameters and number of repeats are reported in Supplementary Table 1.

Substituting Cha and 2,6F_2_-Phe at Phe_cen_ and Phe_ex_ reveals a role for π-electrons of Phe_cen_ for channel gating while sterics and hydrophobicity appear to play a bigger role at Phe_ex_ (Fig. 7B, C, Fig. 7 – Suppl. 1). Neutralization of electrostatics at Phe_cen_ with Cha right-shifts the single-pore G-V by ∼50 mV, while the redistribution of the electrons in 2,6F_2_-Phe left-shifts it by ∼50 mV (Fig. 7C). Conversely, these manipulations have smaller effects at Phe_ex_, as the F214Cha mutant shows an ∼25 mV right-shift of activation and the F214-2,6F_2_-Phe substitution ∼20 mV (Fig. 7B). Similarly, substitutions at Phe_cen_ also affect common-pore gating (Fig. 7E) while those at Phe_ex_ cause only minor alterations (Fig. 7D). Notably, the effect of F418Cha is comparable to that of F418A, suggesting that the electrostatics dominate the interactions of Phe_cen_ with Glu_ex_ during common-pore gating (Fig. 6H, 7E). Finally, we replaced Phe_cen_ with 2,6diMeth-Phe (Fig. 7A), which restricts its interconversion between the ‘up’ and ‘down’ rotamers (Fig. 2E). The kinetics of the common-pore gating process of the F418-2,6diMeth-Phe are accelerated so that they become comparable to those of the single-pore gate (Fig. 7- Suppl. 2A, B). We used the double mutant C212S F418-2,6diMeth-Phe to separate the effects of the non-canonical substitution on the single- and common-pore gating processes (Fig. 7 – Suppl. 2C, D). Using this approach, we found that replacement of Phe_cen_ with the rotameric-restricted isostere causes an almost +40 mV shift in the single-pore gate V_0.5_ and an almost -25 mV shift in the V_0.5_ of the common-pore gating process (Fig. 7C, E). Thus, the conformational rearrangement of Phe_cen_ between the ‘up’ and ‘down’ conformers we identified in the CLC transporters are essential for CLC-0 channel gating, and is necessary to allow the movement of Glu_ex_ in and out of the Cl^−^ permeation pathway. Therefore, our results show that the aromatic slide forms an evolutionarily conserved structural motif that is necessary to enable Glu_ex_ movement in and out of the Cl^−^ pore during the exchange cycle and in channel opening.

## Discussion

Despite extensive structural and functional investigations, the mechanisms underlying the exchange cycle of the CLC transporters and opening of the CLC channels remain poorly understood. These processes are evolutionarily and mechanistically related (Alessio Accardi, 2015; Thomas J. Jentsch & Pusch, 2018; C. Miller, 2006): in both channels and transporters Glu_ex_ moves in and out of the Cl^−^ pathway in a protonation-dependent manner. However, the molecular steps that underlie these rearrangements are unclear. The CLCs exchangers do not follow the ‘ping-pong’ or sequential kinetics that characterize most conventional transporters. In contrast, CLCs simultaneously bind H^+^ and Cl^−^ (Picollo et al., 2012), and substrate movement occurs along two partially congruent translocation pathways (A. Accardi et al., 2005; Zdebik et al., 2008). Yet, how the two substrates bypass each other in the protein remains unknown. The recent structures of the CLC^F^ exchanger and of the CLC-1 Cl^−^ channel suggested that Glu_ex_ might interact with Phe_cen_ (Last et al., 2018; Eunyong Park & MacKinnon, 2018), but the functional implications of these interactions are not clear. Current transport mechanisms for CLC exchangers are not reversible, as they all postulate a step where a protonated and neutral Glu_ex_ displaces Cl^−^ ions bound to S_ext_ and S_cen_ sites to transfer its proton to the internal solution (A. Accardi et al., 2005; Zdebik et al., 2008). This transition is energetically unfavorable (Kuang et al., 2007), and would result in highly asymmetric transport rates. However, the CLC exchangers function with comparable efficiency in both directions (De Stefano et al., 2013; L. Leisle et al., 2011; Matulef & Maduke, 2005), arguing against the existence of an asymmetric rate-limiting step. Here, a combination of MD simulations with biochemical and electrophysiological experiments suggest that Glu_ex_ takes different routes depending on its protonation state.

### Formation of an aromatic slide is essential to Glu_ex_ movement

Our data suggest that while the deprotonated and negatively charged Glu_ex_ moves through the Cl^−^ pathway, the protonated and neutral Glu^0^_ex_ interacts with two highly conserved phenylalanines, Phe_cen_ and Phe_ex_. These residues can form an aromatic slide that enables movement of protons bound to Glu_ex_ in and out of the pathway, connecting the central binding site and the extracellular solution. In the available structures of CLC channels and exchangers (A. Accardi et al., 2006; A. Accardi et al., 2005; Basilio et al., 2014; Dutzler et al., 2002; Dutzler et al., 2003; Feng et al., 2010; Jayaram, Robertson, Wu, Williams, & Miller, 2011; Last et al., 2018; Lobet & Dutzler, 2006; E. Park et al., 2017; Eunyong Park & MacKinnon, 2018; Wang et al., 2019), Phe_cen_ is in the ‘up’ rotamer suggesting this conformation is likely the most stable. However, our combined simulation and functional data suggests that Phecen can adopt two distinct rotamers around its χ1 angle (Fig. 2), and that this rearrangement determines the energy barrier for Cl^−^ movement within the pore (Fig. 2, Fig. 2 - Suppl. 2) and enables Glu^0^_ex_ to reach or leave S_cen_ via the aromatic slide pathway (Fig. 3). Further, we show that the aromatic slide residues play an evolutionarily conserved key role in the CLC exchange cycle and in CLC channel gating. Our conventional mutagenesis underscores the conserved importance of Phe_cen_ and Phe_ex_ in transporter and channel gating: alanine substitutions at these positions impair the absolute transport rate, the voltage dependence and the exchange stoichiometry of the transporters (Fig. 4, 5), and nearly abolish the voltage dependence of channel gating (Fig. 6). We used atomic scale mutagenesis to test the prediction of our MD simulations that formation of the aromatic slide entails a rotation of the side chain of Phe_cen_ around its C_α_-C_β_ bond. When we replace Phe_cen_ with 2,6diMet-Phe, a Phe derivative that specifically constrains this rotational rearrangement, we find that opening of the single- and common-pore gates in the CLC-0 channel are severely impaired (Fig. 7, Fig. 7 – Suppl. 1, 2). Thus, the aromatic slide forms an evolutionarily conserved structural motif that enables movement of a protonated Glu^0^_ex_ in and out of the pathway in both CLC channels and transporters, consistent with the finding that some CLC channels mediate some residual H^+^ transport (Lísal & Maduke, 2008; Picollo & Pusch, 2005). The low expression of the CLC-5 and -7 transporters encoding the non-canonical amino acid prevented a similar test in these homologues.

Our results also suggest that Phe_ex_ and Phe_cen_ interact differently with Glu_ex_. The aromatic ring of Phe_ex_ plays a structural role in helping position Glu_ex_ where it can interact with Phe_cen_ in the ‘up’ conformation (Fig. 3). Indeed, removal of the aromatic ring of Phe_ex_ severely affects voltage dependent gating of CLC channels and transporters (Fig. 5, 6), while selective manipulations of its electrostatic properties have only minor effects on channel gating (Fig. 7B, D). In contrast, the electrostatic properties of the aromatic ring of Phe_cen_ are essential determinants for Glu_ex_ movement in the CLC-0 channel: elimination of the π-electrons of Phe_cen_ favors the closed state of the single-pore gate (Fig. 7C), while their re-localization to the proximal edge promotes opening (Fig. 7C). These findings are consistent with the location of Phe_cen_ within the core of the protein, and with our MD simulations suggesting that Glu_ex_ forms a π-dipole interaction with the aromatic ring of Phe_cen_ (Fig. 3). Indeed, the interaction between the buried aromatic Phe_cen_ and Glu_ex_ could account for the observed shifts in the pKa of Glu_ex_ (Robinson, Majumdar, Schlessman, & García-Moreno E, 2017) in CLC channels and transporters to keep Glu_ex_ protonated during its transition through the protein (Hanke & Miller, 1983; Niemeyer, Cid, Yusef, Briones, & Sepúlveda, 2009; Picollo, Malvezzi, & Accardi, 2010; Picollo et al., 2012). Finally, our results are in harmony with the recent proposals that Glu_ex_ can adopt a conformation where it directly interacts with Phe_cen_ in a fluoride selective CLC antiporter (Last et al., 2018) and in the CLC-1 channel (Eunyong Park & MacKinnon, 2018) (Fig. 4 Suppl. 2C).

### A mechanism for Cl^−^/H^+^ exchange

Our finding that two Phe residues form an evolutionarily conserved secondary pathway that enables movement of the protonated Glu_ex_ in and out of the ion transport pathway allows us to propose a 7-state mechanism for the CLC transporters that explains the stoichiometry of 2 Cl^−^:1 H^+^, is fully reversible, and accounts for previous results. For simplicity of representation, we consider that an ion in the S_ext_* site is out of the pathway and as such, in our scheme we do not show ions bound to this site. As a starting configuration, we consider a state where Glu_ex_ and Glu_in_ are de-protonated, Glu_ex_ occupies S_cen_, Phe_cen_ is in the ‘up’ position and no Cl^−^ ions are bound to the pathway (Fig. 8, I). After a Cl^−^ ion binds to S_int_ (Fig. 8, II), the opening of the intracellular gate, formed by Ser_cen_ and Tyr_cen_ (Basilio et al., 2014), allows the ion to move into S_cen_, displacing Glu_ex_ to S_ext_ (Fig. 8, III). The binding of a second ion favors the protonation of Glu_ex_, allowing the Cl^−^ ions to simultaneously occupy S_cen_ and S_ext_, accompanied by the displacement of Glu^0^_ex_ out of the pathway and the closure of the internal gate (Fig. 8, IV). The protoned Glu_ex_ diffuses toward the aromatic slide and interact with Phe_ex_ (Fig. 8, V). Movement of Glu^0^_ex_ along the aromatic slide allows the interdependent rearrangement of Phe_cen_ to the ‘down’ state and release of a Cl^−^ ion, i.e. release from S_ext_ to the extracellular milieu and the transfer of the second ion from S_cen_ to S_ext_ (Fig. 8, VI). This conformation, where Glu^0^_ex_ interacts with Phe_cen_ on the side of the Cl^−^ pathway and S_ext_ is occupied, favors the formation of a water wire (Fig. 8, VI), which enables proton transfer from Glu^0^_ex_ to Glu_in_ (Fig. 8, VII). Deprotonation of Glu_ex_ allows it to move into S_cen_, favoring the release of the second Cl^−^ from S_ext_ to the outside, and of the proton from Glu^0^_in_ to the intracellular solution, returning to the starting configuration (Fig. 8, I). In sum, we propose that the Cl^−^ and H^+^ ions bypass each other while moving in opposite directions through a CLC transporter by taking physically distinct routes: the Cl^−^ ions move through the anion selective pore while the H^+^ moves along a pathway comprised of a water-wire and the aromatic slide (Fig. 8B). The distinct routes for Cl^−^ and H^+^ allow for a complete reversibility of the cycle.

**Figure 8.**
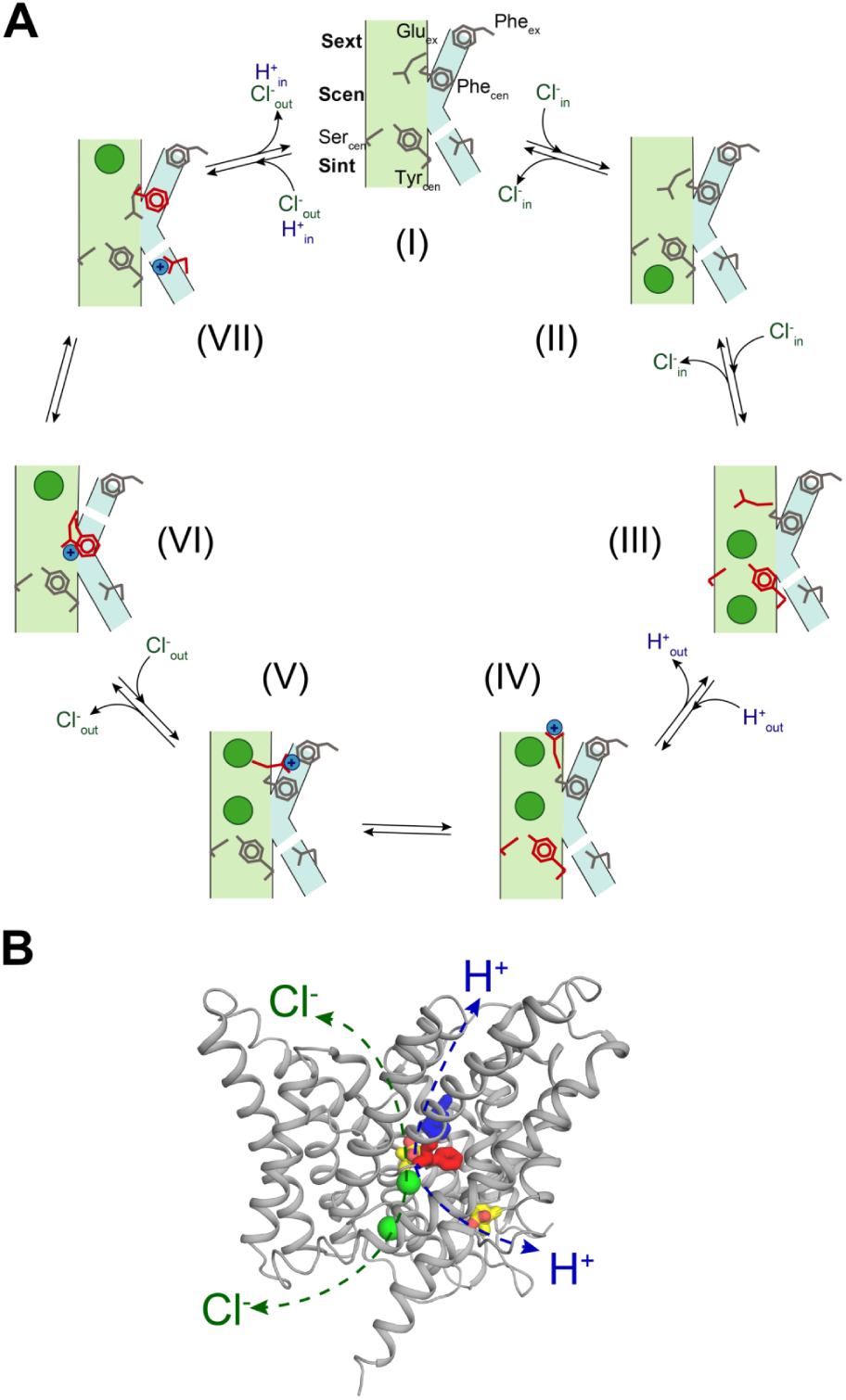
Proposed mechanism for the 2 Cl-: 1 H+ CLC exchangers. (**A**) Schematic representation of the 2 Cl^−^: 1 H^+^ CLC exchange cycle. The Cl^−^ ions are shown as green circles and H^+^ as blue circles. The Cl^−^ and H^+^ pathways are shown in pale green and cyan respectively. For clarity, the residues that undergo conformational rearrangements in each step are highlighted in red. The H^+^ pathway is shown as discontinuous when it is in a non-conductive conformation. (I) Apo- and occluded-state of the transporter with Glu_ex_ in S_cen_ and Phe_cen_ in the up rotamer. (II) An intracellular Cl^−^ ion binds to S_int_. (III) The inner gate opens, a second intracellular Cl^−^ binds so that both S_int_ and S_cen_ are occupied and Glu_ex_ moves to S_ext_. (IV) The Cl^−^ ions move to S_cen_ and S_ext_, the inner gate closes, Glu_ex_ moves out of the pathway and becomes protonated. (V) Glu^0^_ex_ interacts with Phe_ex_. (VI) Glu^0^_ex_ interacts with Phe_cen_ which rotates to the down conformation enabling the formation of a water wire connecting Glu^0^_ex_ and Glu_in_; one Cl^−^ ion is released from S_ext_ and the other one moves from S_cen_ to S_ext_. (VII) The H^+^ is transferred to Glu_in_ favoring movement of Glu_ex_ into S_cen_ and the Cl^−^ ion in S_ext_ is released to the extracellular solution (I). (**B**) Physically distinct Cl^−^ and H^+^ pathways are indicated in CLC-ec1 WT structure (Glu_ex_ and Glu_in_ are shown in yellow, Phe_ex_ in blue, Phe_cen_ in red, Cl^−^ ions as green spheres). Both pathways converge at Glu_ex_.

Remarkably, our findings show that the role of the aromatic slide residues Phe_cen_ and Phe_ex_ is evolutionarily conserved between CLC exchangers and channels. We propose that the aromatic slide in CLC-0 could provide a conserved pathway that enables the residual H^+^ transport associated with the common-pore gating process of the CLC-0 channel to occur (Lísal & Maduke, 2008). Thus, our proposed exchange mechanism for the CLC transporters also captures the key rearrangements that underlie gating of the CLC channels. In this framework, rapid ion conduction would be enabled by the disruption of the intracellular gate, a hypothesis justified by the widened intracellular vestibule seen in the recent structures of the bCLC-K and hCLC-1 channels (E. Park et al., 2017; Eunyong Park & MacKinnon, 2018; Wang et al., 2019).

## Materials and Methods

### Molecular system

The molecular systems, based on the CLC-ec1 crystal structure (PDB: 1OTS) (Dutzler et al., 2003), were assembled using the CHARMM-GUI web service (Jo, Kim, Iyer, & Im, 2008). As it was shown that the monomeric form of CLC-ec1 is functional (Robertson, Kolmakova-Partensky, & Miller, 2010), a single subunit was used. The x-ray structure includes residues 18 to 458. Residues 18 to 31 are normally interacting with the opposite subunit and were removed here to prevent any unexpected interaction with the pore. All residues are in their natural protonation state at pH 7 except Glu148, which is tested in both its unprotonated and protonated forms. The orthorhombic periodic simulation cell contains a ClC-ec1 monomer inserted in a bilayer composed of 262 dimystoylphosphatidylcholine (DMPC) lipids, about 13 600 TIP3P water molecules, and potassium/chloride ions to neutralize the systems at a concentration of about 150 mM KCl.

### Potential function and simulations

All simulations were run using the CHARMM simulation software version c36 (Brooks et al., 2009) using the CHARMM36 force field (Best et al., 2012). Simulations were performed in an isothermal−isobaric ensemble with a pressure of 1 atm and a temperature of 323 K. Particle-mesh Ewald method (Essmann et al., 1995) was applied for the calculation of electrostatic interactions with a grid spacing of 1 Å. The cutoff distance for van der Waals interactions was taken at 12 Å with a switching function starting at 10 Å. Time step for the integration of the motion was set to 2 fs. The membrane systems were equilibrated following a standard protocol offered by CHARMM-GUI (Wu et al., 2014). After equilibration, a simulation of 100 ns was run for each molecular system prior to the free energy calculations.

### Potential of mean force calculations

The potential of mean forces (PMF) were calculated using the self-learning adaptive umbrella sampling method (Wojtas-Niziurski, Meng, Roux, & Bernèche, 2013). In the case of PMFs describing the displacement of one or two ions, the reaction coordinate is the position of a given ion in the pore, as projected on the normal to the membrane (Z axis). The reference point is the center of mass of the backbone of residues 107, 356, 357, and 445. Independent simulations of 500 ps were performed every 0.5 Å along the reaction coordinate using a biasing harmonic potential of 20 kcal/mol•Å^2^. In the case of PMFs describing the rotation of the F357 side chain, the reaction coordinate is the *χ*1 angle. Simulations were performed every 10 degrees using a biasing harmonic potential of 20 kcal/mol•rad^2^. In all cases the first 50 ps of every independent window simulation was considered as equilibration and thus excluded from the analysis. The independent simulations were combined and unbiased using the weighted histogram analysis method (WHAM) (Kumar, Rosenberg, Bouzida, Swendsen, & Kollman, 1992).

### Protein purification and liposome reconstitution

Expression and purification of wild-type and mutant CLC-ec1 were performed according to published protocols (A. Accardi, Lobet, Williams, Miller, & Dutzler, 2006; A. Accardi & Miller, 2004; Basilio et al., 2014; Picollo et al., 2009; Picollo et al., 2012). Purified proteins were reconstituted into liposomes as described(Basilio et al., 2014).

### Cl^−^ and H^+^ flux recordings for CLC-ec1 and variants

Cl^−^ and H^+^ fluxes of CLC-ec1 wild-type and mutant proteins reconstituted into proteoliposomes were recorded simultaneously and the coupling stoichiometry was determined as described (Basilio et al., 2014).

### Isothermal titration calorimetry (ITC)

Cl^−^ binding affinity was determined for purified CLC-ec1 F357A as described (Basilio et al., 2014; Picollo et al., 2009) using a nanoITC instrument (TA Instruments). For these experiments, the final purification step of the protein was purified over a gel filtration column pre-equilibrated in 100 mM Na-K-Tartrate, 20 mM Hepes, 50 μM DMNG, pH 7.5 (Buffer B0) and concentrated to 50-195 μM. The injection syringe was filled with buffer B0 with 50 mM KCl added, to achieve final Molar Ratios of 70-100. Each experiment consisted of 30-48 injections of 1 μl of the ligand solution at 3-4 min intervals into the experimental chamber kept under constant stirring at 350 rpm and at 25.0±0.1 °C. All solutions were filtered and degassed prior to use. The ITC data was fit to a single site Wiseman isotherm using the NanoAnalyze program from TA instruments.

### In vitro cRNA transcription

RNAs for all CLC-0, hCLC-7, hCLC-5 and mOstm1 wild-type and mutant constructs were transcribed from a pTLN vector using the mMessage mMachine SP6 Kit (Thermo Fisher Scientific, Grand Island, NY) (L. Leisle et al., 2011; Pusch, Ludewig, Rehfeldt, & Jentsch, 1995; Steinmeyer, Schwappach, Bens, Vandewalle, & Jentsch, 1995). For all experiments in this paper a plasma-membrane localized version of hCLC-7 has been used as ‘wild-type’ which has been published earlier termed as CLC-7^PM^ (L. Leisle et al., 2011). For final purification of cRNA the RNeasy Mini Kit (Quiagen, Hilden, Germany) was employed. RNA concentrations were determined by absorbance measurements at 260 nm and quality was confirmed on a 1% agarose gel.

### tRNA misacylation

For nonsense suppression of CLC-0 TAG mutants (F214X and F418X) in Xenopus laevis oocytes THG73 and PylT tRNAs have been employed. THG73 was transcribed, folded and misacylated as previously described (Lilia Leisle et al., 2016). PylT was synthetized by Integrated DNA Technologies, Inc. (Coralville, IA, USA), folded and misacylated as described(Infield, Lueck, Galpin, Galles, & Ahern, 2018). Phe-, Cha-, 2,6F2-Phe- and 2,6diMethPhe-pdCpA substrates were synthesized according to published procedures (Infield et al., 2018). L-Cha was purchased from ChemImpex (Wood Dale, IL, USA; catalogue number: 11696) and BACHEM (USA; catalogue number: F-2500.0001), L-2,6-difluoro Phe from ChemImpex (catalogue number: 24171) and L-2,6-dimethyl Phe from Enamine (Monmouth Junction, NJ, USA; catalogue number: EN300-393063).

### Protein expression in Xenopus laevis oocytes and two electrode voltage clamp (TEVC) recordings

Xenopus leavis oocytes were purchased from Ecocyte Bio Science (Austin, TX, USA) and Xenoocyte (Dexter, Michigan, USA) or kindly provided by Dr. Pablo Artigas (Texas Tech University, USA, protocol # 11024). For conventional CLC expression, following injection and expression conditions have been used: for CLC-7 (WT, F301A or F514A) and Ostm1, 25 ng of each cRNA were injected per oocytes and currents were recorded after ∼60-80 h; for CLC-5 (WT, F255A or F445A), 50 ng cRNA were injected per oocyte and currents were recorded ∼48-72 h; for CLC-0 (WT, F214A or F418A), 0.1-5 ng cRNA were injected and currents were measured ∼6-24 h after injection. For nonsense suppression of CLC-0 constructs (F214X, F418X, C212S F418X), cRNA and misacylated tRNA were coinjected (up to 25 ng of cRNA and up to 250 ng of tRNA per oocyte) and currents were recorded 6-24 h after injection.

TEVC was performed as described (Picollo et al., 2009). In brief, voltage-clamped chloride currents were recorded in ND96 solution (in mM: 96 NaCl, 2 KCl, 1.8 CaCl2, 1 MgCl2, 5 HEPES, pH 7.5) using an OC-725C voltage clamp amplifier (Warner Instruments, Hamden, CT). The data were acquired with the GePulse software (M. Pusch, Istituto di Biofisica, Genova) and analyzed using Ana (M. Pusch, Istituto di Biofisica, Genova) and Prism (GraphPad, San Diego, CA, USA). Standard voltage-clamp protocols have been applied for the three CLC proteins, the holding potential was constant at -30 mV. For CLC-0 two different recording protocols have been used to distinguish single-pore from common-pore gating. During the single-pore gating protocol the voltage was stepped to +80 mV for 50 ms and then a variable voltage from -160 mV to +80 mV increasing in 20 mV steps was applied for 200 ms, followed by a 50 ms pulse at -120 mV for tail current analysis. For CLC-0 common-pore gating, 7 s voltage steps from +20 mV to -140 mV have been applied in -20 mV increments followed by a 2.5 s +60 mV post pulse for tail current analysis. For CLC-5, a simple voltage step protocol was applied: 400 ms steps from -80 to + 80 mV in 20 mV increments. For CLC-7/Ostm1, voltage was clamped at variable values from -80 to +90 mV in 10 mV steps for 2 s, followed by a 0.5 s post pulse at -80 mV for tail current analysis.

To estimate the voltage dependence of CLC-7/Ostm1 and CLC-0 gating, tail current analysis was performed and data were fit to a Boltzman function of the form:

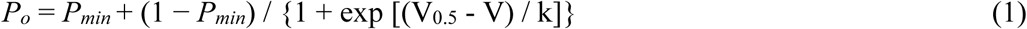

where *P_o_* is the open probability as a function of voltage and is assumed to reach a value of unity at full activation. *P_min_* is the residual open probability independent of voltage. V_0.5_ is the voltage at which 50% activation occurs, and k is the slope factor (k=R*T/(z*F) with R as universal gas constant, T as temperature in K, F as Faraday constant, and z as the gating charge).

For analysis of the activation kinetics of CLC-7/Ostm1 and its variants, activating voltage pulses (from +20 to +90 mV) were fit to a bi-exponential function of the following form:

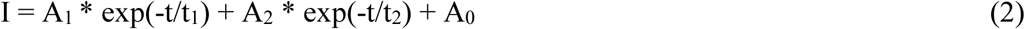

where I is the current as a function of time; A_1_, A_2_ and A_0_ are fractional amplitudes obtained by normalizing to the total current. While A_1_ and A_2_ are time-dependent components, A_0_ is time-independent. τ_1_ and τ_2_ are the corresponding time constants.

### Statistical analysis

All values are presented as mean ± s.e.m. To determine statistical significance Student’s t-test (two-tailed distribution; two-sample equal variance) was performed. The threshold for significance was set to *P*=0.05.

## Acknowledgements

The authors wish to thank members of the Accardi lab for helpful discussions. This work was supported by NIH/NIGMS grant 1R01GM128420 (to A.A.), Swiss National Science Foundation SNF Professorship No PP00P3_139205 (to S.B.) and NIH/NINDS grant NS104617 to (C.A.A.)

## Author contributions

Y.X. and S.B. designed, carried out and analyzed all MD simulations; L.L. and A.A. designed, carried out and analyzed all electrophysiological experiments; E.F., M.V. and A.A. designed, carried out and analyzed all biochemical experiments; J.G. and C.A.A. designed and generated all non-canonical amino-acids; A.A., S.B. and C.A.A. designed and oversaw the research; L.L., Y.X., S.B. and A.A. wrote the initial manuscript, all authors edited the manuscript.

## Data availability

Data supporting the findings of this manuscript are available from the corresponding authors upon request.

## SUPPLEMENTARY INFORMATION

**Figure 2 - Supplement 1.**
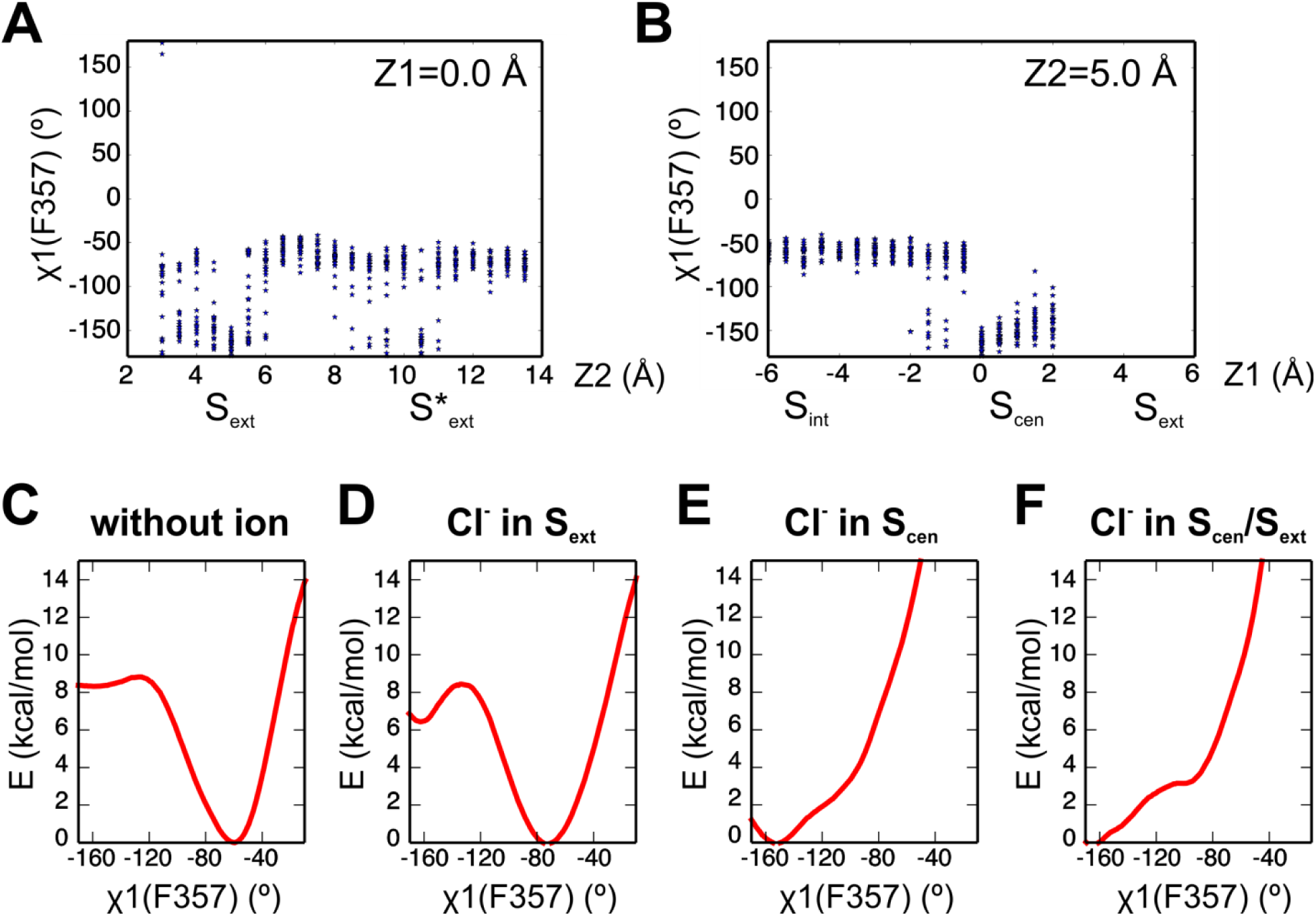
Ion binding influences the conformation of the F357 side chain. (**A,B**) The conformational sampling underlying the PMF of Fig. 2A reveals large fluctuations of F357 *χ*1 angle, notably when a Cl^−^ visits the S_cen_ binding site. The S_cen_/S_ext_ doubly occupied state correspond to the coordinate (Z1, Z2) = (0.0, 5.0). The distribution of F357 *χ*1 values is plotted for each simulation window along the axes Z1= 0.0 Å (A) and Z2= 5.0 Å (B). (**C-F**) The PMF along F357 *χ*1 angle is shown for different occupancy states: in absence of ion (**C**), with a Cl^−^ in S_ext_ (**D**) or S_cen_ (**E**), and with Cl^−^ in both S_cen_ and S_ext_ (**F**).

**Figure 2 - Supplement 2.**
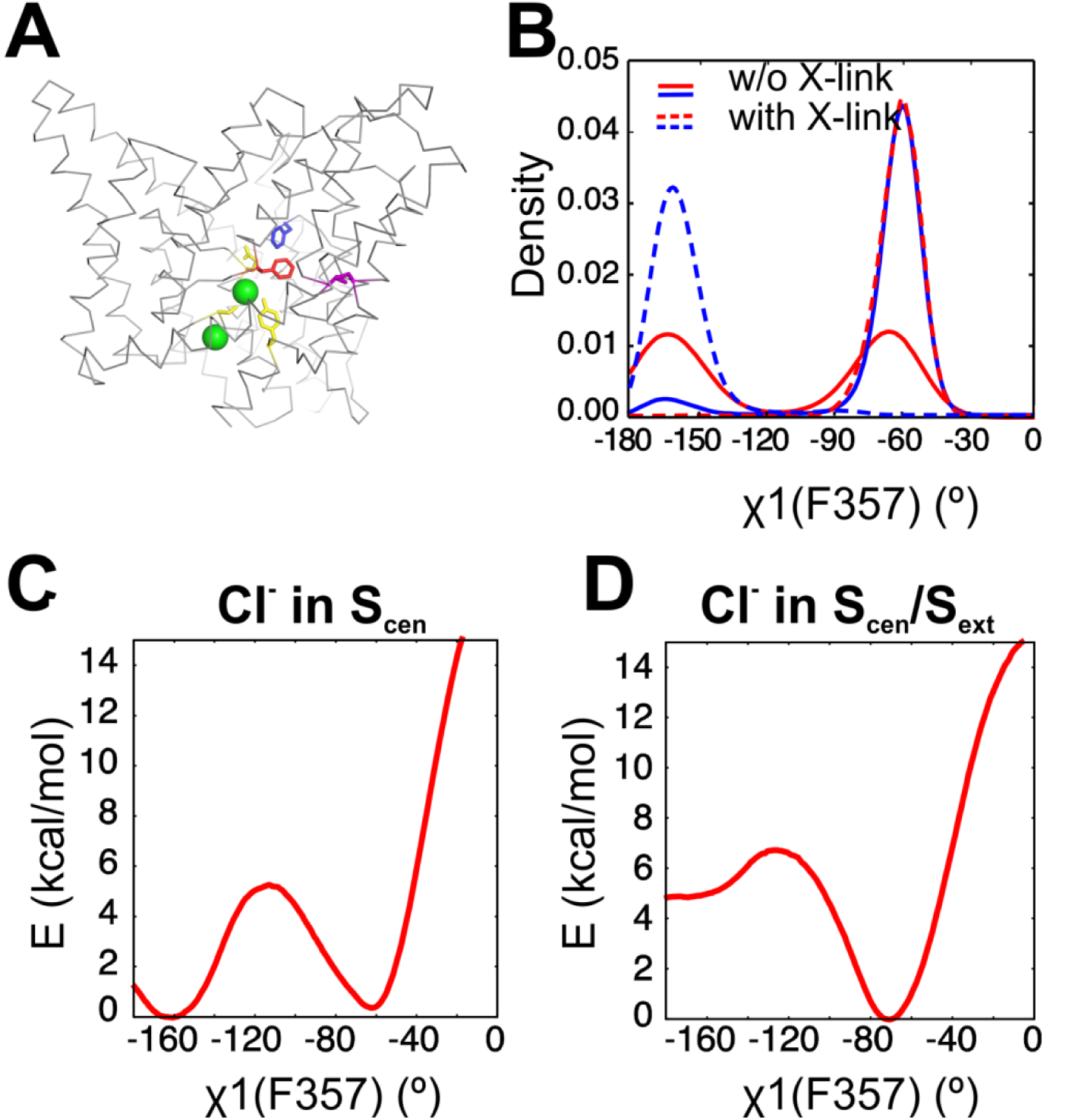
A crosslink known to reduce transport activity impedes the reorientation of the F357 side chain. (**A**) Molecular graphic shows the position of the engineered crosslink within a CLC-ec1 monomer with the A399C-A432C crosslink in magenta sticks, the Cl-coordinating residues (S107, E148 and Y445) in yellow sticks, Phe_cen_ (F357) and Phext (F190) respectively as red and blue sticks, Cl^−^ ions as green spheres. (**B**) The distribution of F357 *χ*1 angle is shown for simulations in absence and presence of the engineered crosslink. Two independent simulations are shown for each case. (**C, D**) The PMFs along F357 *χ*1 angle shown in Figure 2 – Supplement 1 E, F are repeated here in presence of the crosslink, with a Cl^−^ in S_cen_ (**C**) and Cl^−^ in both S_cen_ and S_ext_ (**D**). The Cys bridge increases the free energy barrier between the F357 down (*χ*1= -70°) and up (*χ*1= -160°) conformations.

**Figure 3 - Supplement 1.**
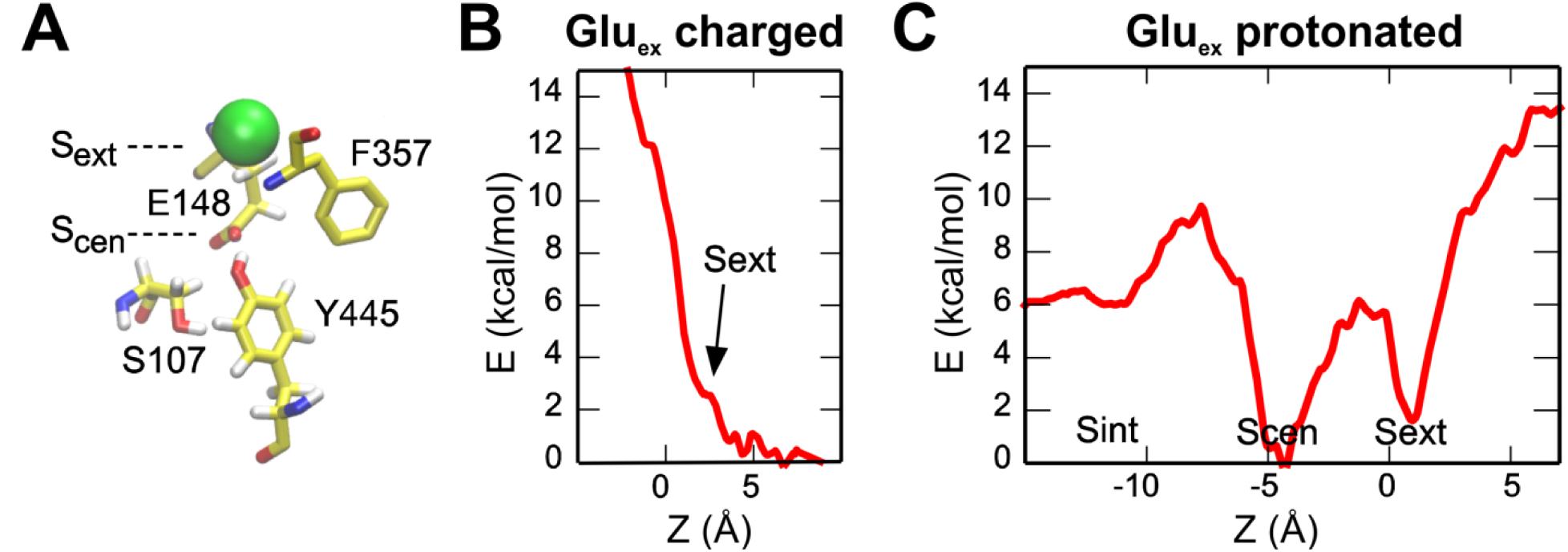
Binding of a Cl^−^ to S_ext_ and S_cen_ with Glu_ex_ in the pore. (**A**) The molecular representation shows the initial state used for the PMF calculations. (**B, C**) The PMFs describe the entry of a Cl^−^ into the pore when Glu_ex_ is charged (non-protonated) (**B**) or neutral (protonated) (**C**). The reaction coordinate is defined as in Figure 2. S_cen_ and S_ext_ binding sites are accessible only when Glu_ex_ is protonated, in agreement with the idea that Cl^−^ and H^+^ binding happens simultaneously. The PMFs also show that S_int_ presents a lower binding affinity than S_cen_ and S_ext_.

**Figure 4 - Supplement 1.**
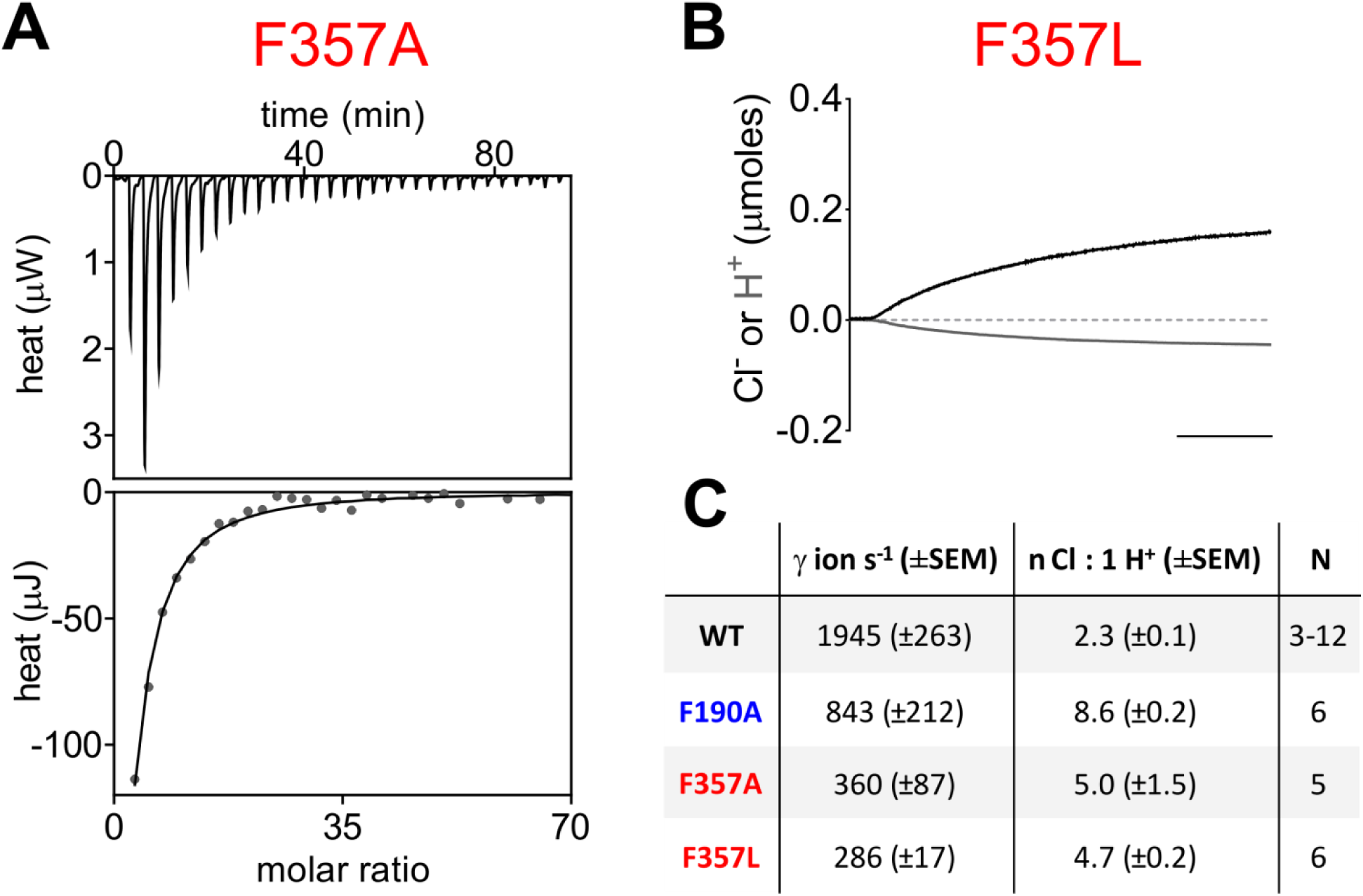
Effect of mutations at F357 on ion binding and transport in the CLC-ec1 exchanger. **(A)** Thermograms of Cl^−^ binding to CLC-ec1 F357A. Top graphs show the heats released upon ion binding. Bottom graphs show the integrated heats (circles), and the solid line is the fit to a single-site isotherm with K_D_=0.67±0.02 mM; ΔH= -5.1±0.3 kcal Mol^−1^, TΔS= -0.8±0.3 kcal Mol^−1^, and n=1 binding site, N=2 independent repeats. **(B)** Representative time course of Cl^−^ (black) and H^+^ (gray) transport recordings of purified CLC-ec1 F357L reconstituted into liposomes. *Scale bar*, 25 s. **(C)** Summary table of transport rate and coupling stoichiometry for WT and mutant CLC-ec1. Numbers (N) of independent experiments are indicated.

**Figure 4 - Supplement 2.**
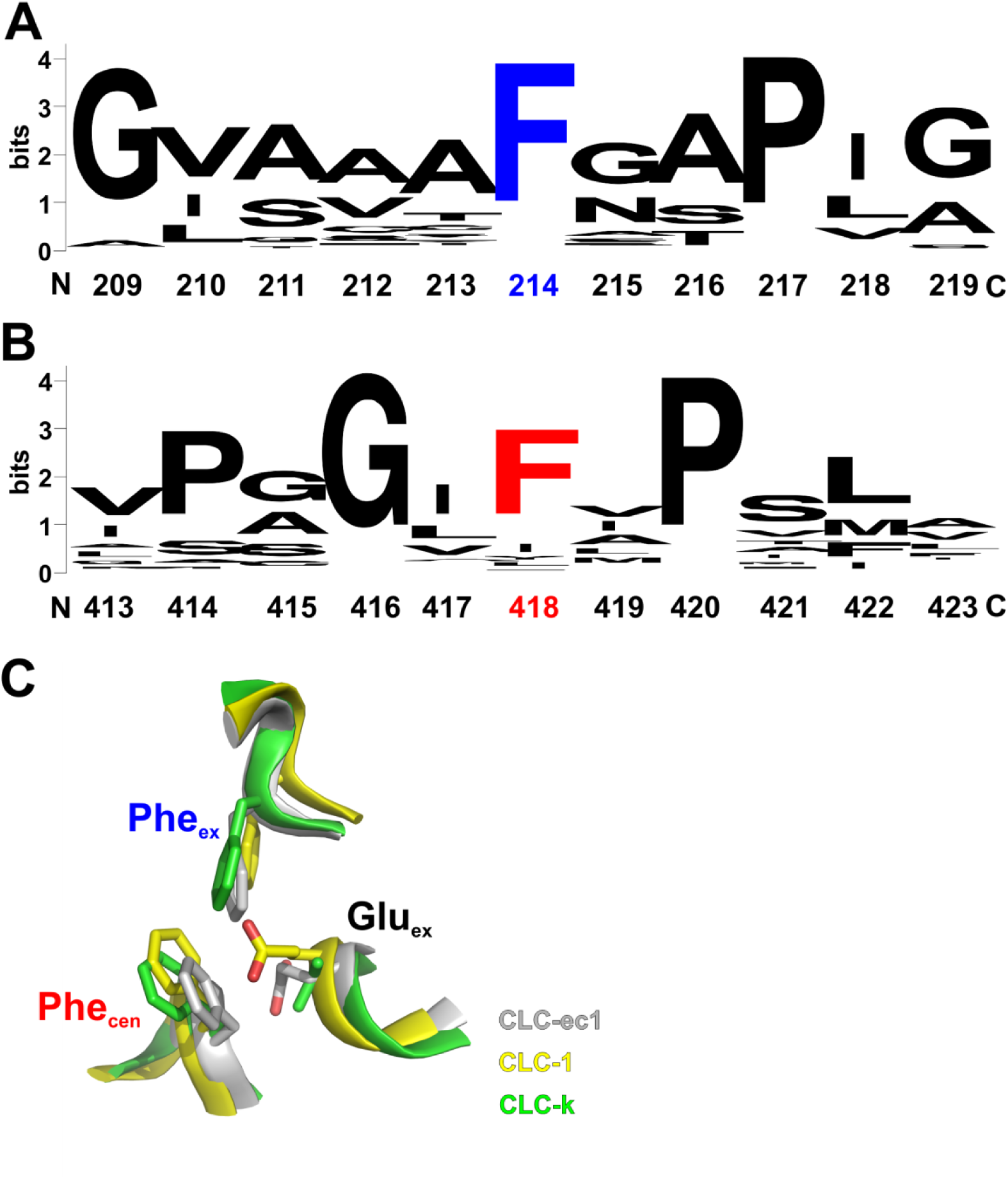
Conservation of Phe_ex_ and Phe_cen_ within the CLC family. **Phe_ex_** (**A**) and Phe_cen_ (**B**) show a high degree of conservation across the CLC family: 94% and 76%, respectively, as calculated from an alignment of 2,200 CLC sequences. Logos were created using the Weblogo server (https://weblogo.berkeley.edu/logo.cgi). (**C**) Superposition of the extracellular region of the Cl^−^ and H^+^ pathways of the CLC-ec1 exchanger (grey; PDBID: 1OTS), the CLC-1 channel (yellow; PDBID: 6COY) and of the CLC-k channel (green; PDBID: 5TQQ). Phe_ex_, Phe_cen_ and Glu_ex_ are shown as sticks with the oxygen atoms in red; parts of the backbone are shown as ribbons. Note that in the CLC-k channel Glu_ex_ is replaced by a valine.

**Figure 5 - Supplement 1.**
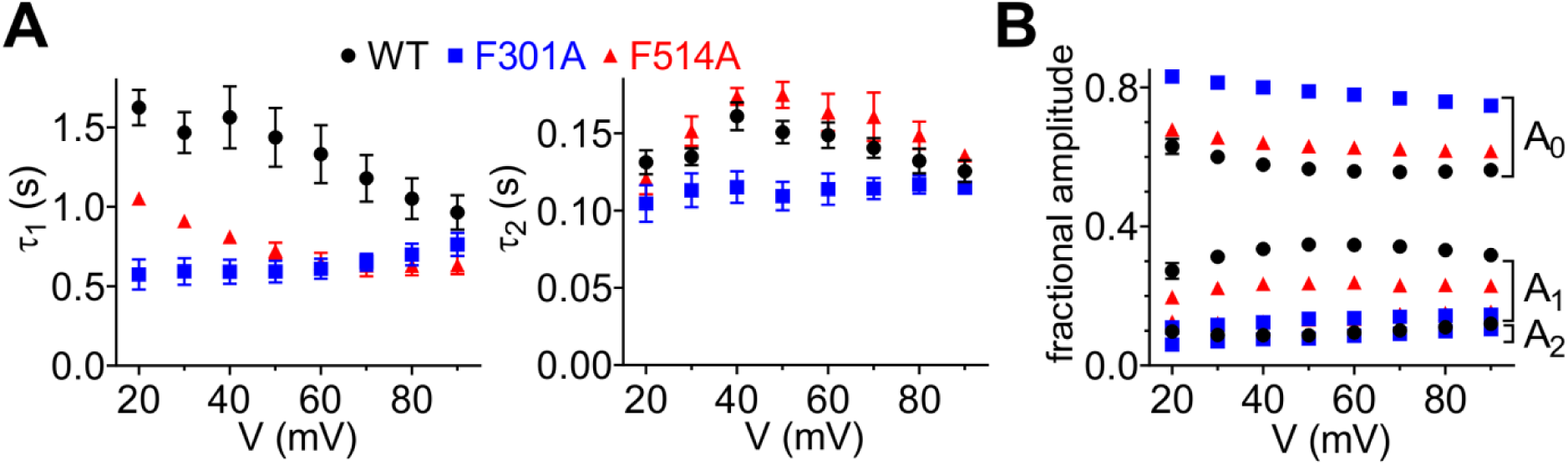
Phe_cen_ (F514) and Phe_ex_ (F301) impact activation kinetics of CLC-7 exchanger. The time courses of current activation of CLC-7 WT (N=8), F301A (N=8) and F514A (N=8) at voltages between +20 and +90 mV were fit to a biexponential function (see *Methods*, Equation 2). **(A)** The slow (τ_1_, left) and fast (τ_2_, right) time constants of CLC-7 WT (black circles), F301A (blue squares) and F514A (red triangles) are plotted as a function of voltage. F301A strongly diminishes voltage dependence of both activation time constants (τ_1_, τ_2_; at every tested voltage *P*<0.05). F514A accelerates the slow time component τ_1_ (at every tested voltage *P*<0.005). **(B)** The fractional amplitudes A_0_, A_1_ and A_2_ for CLC-7 WT (black circles), F301A (blue squares) and F514A (red triangles) are plotted as a function of voltage. F301A increases the time-independent fractional amplitude A_0_ and decreases the contribution of time-dependent fractional amplitude A_1_ (at every voltage tested *P*<0.005). Also F514A increases A_0_ and decreases A_1_ (at every voltage tested *P*<0.005). Values are reported as mean ± S.E.M, error bars are not shown where they are smaller than the symbol size.

**Figure 5 - Supplement 2.**
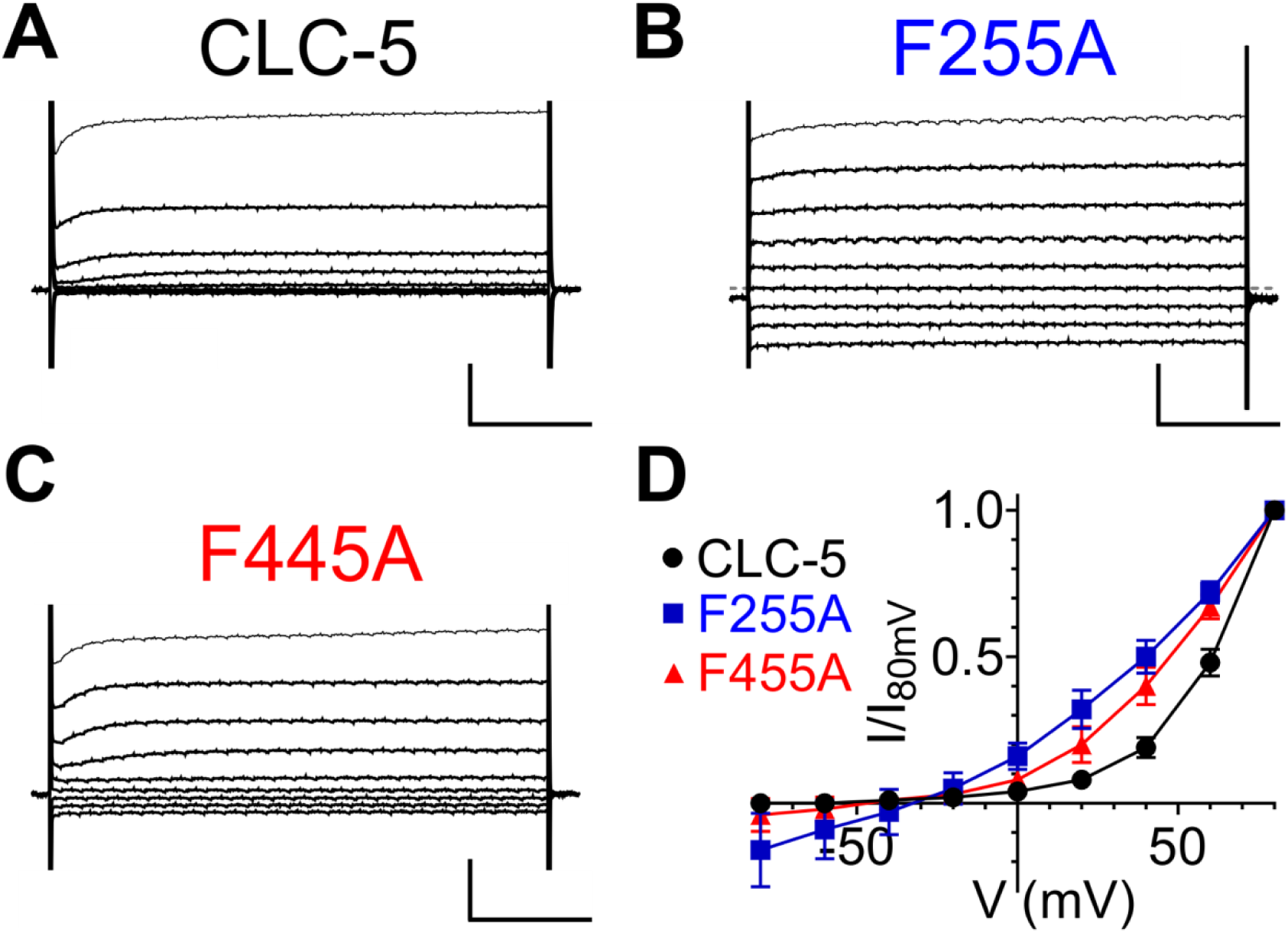
Phe_cen_ (F445) and Phe_ex_ (F255) regulate the voltage dependence of the CLC-5 exchanger. **(A-C)** TEVC current recordings of CLC-5 WT **(A)**, F255A **(B)** and F445A **(C)**. For voltage clamp protocol see *Methods* section. *Horizontal scale bar*, 100 ms; *vertical scale bar*, 0.5 μA. **(D)** Normalized steady state I-V relationship for CLC-5 WT (black circles, N=25), F255A (blue squares, N=11) and F455A (red triangles, N=22). Values are reported as mean ± S.E.M, error bars are not shown where they are smaller than the symbol size.

**Figure 7 - Supplement 1.**
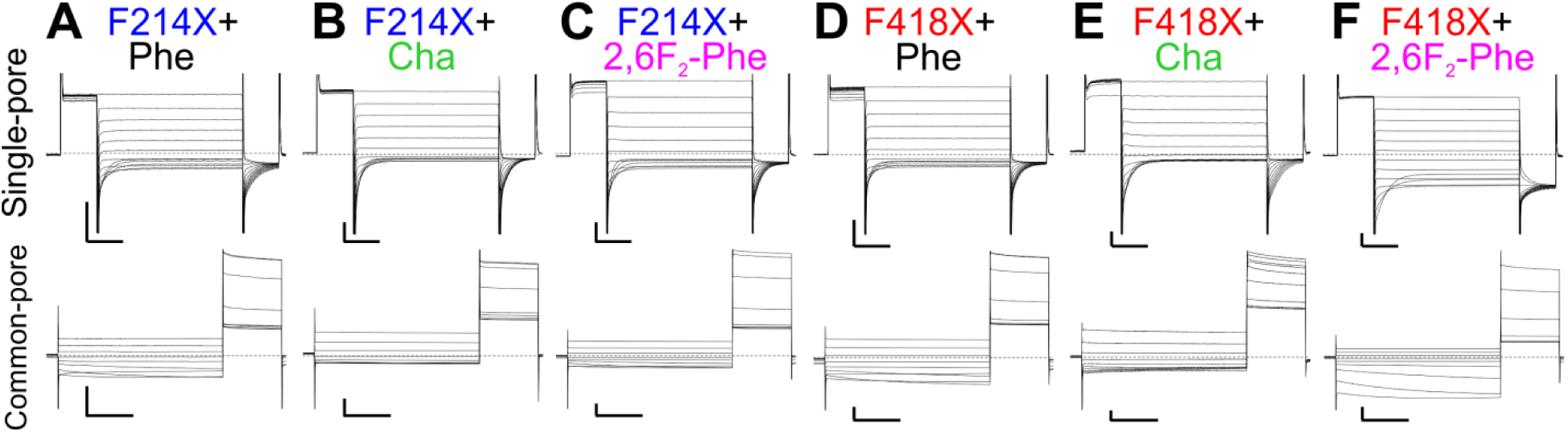
Effects of charge redistribution at Phe_ex_ and Phe_cen_ on the single- and common-pore gating processes of CLC-0. **(A-F)** Representative TEVC currents evoked with single- (top) and common-pore (bottom) gating protocols of the following CLC-0 variants: Phe_ex_ (F214X) replacements with Phe (**A**), Cha (**B**) or 2,6F_2_-Phe (**C**); Phe_cen_ (F418X) substitutions with Phe (**D**), Cha (**E**) or 2,6F_2_-Phe (**F**). Horizontal scale bar, 50 ms (single-pore gate) and 2 s (common-pore gate), respectively; vertical scale bar, 2 μA (single-pore gate) and 1 μA (common-pore gate), respectively.

**Figure 7 - Supplement 2.**
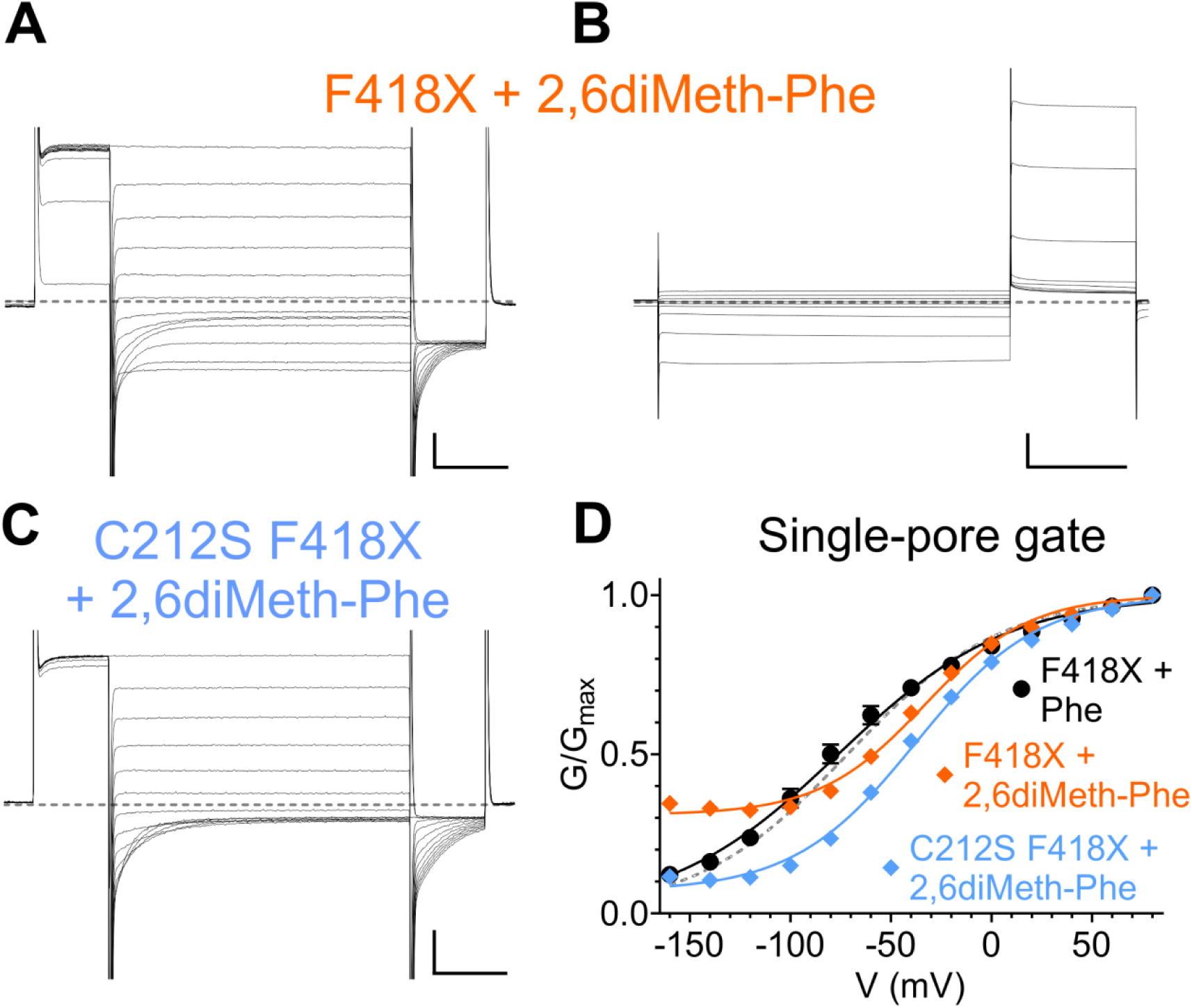
Effects of CLC-0 F418X-2,6diMeth-Phe on single- and common pore gating processes. **(A-B)** Representative TEVC currents of CLC-0 F418X+2,6diMeth-Phe currents evoked with single-(**A**) and common-pore (**B**) gating protocols. **(C)** Representative TEVC currents of CLC-0 C212S F418X+2,6diMeth-Phe evoked with the single-pore gating protocol. *Horizontal scale bars*, 50 ms (single-pore gate) and 2 s (common-pore gate), respectively; *vertical scale bars*, 1 μA (single-pore gate) and 2 μA (common-pore gate), respectively. **(D)** Normalized G-V relationships of the single-pore gating processes of CLC-0 F418X+Phe (black circles; from Fig. 7), F418X+2,6diMeth-Phe (orange diamonds) and C212S F418X-2,6diMeth-Phe (cyan diamonds; from Fig. 7). Solid lines represent fits to a Boltzmann function with an offset (see *Methods*, Equation 1). WT G-V curves (from Fig. 6) are shown as gray dashed lines for reference. Values are reported as mean ± S.E.M, error bars are not shown where they are smaller than the symbol size. Values for the fit parameters and number of repeats are reported in Supplementary Table 1.

**Figure 7 - Supplement 3.**
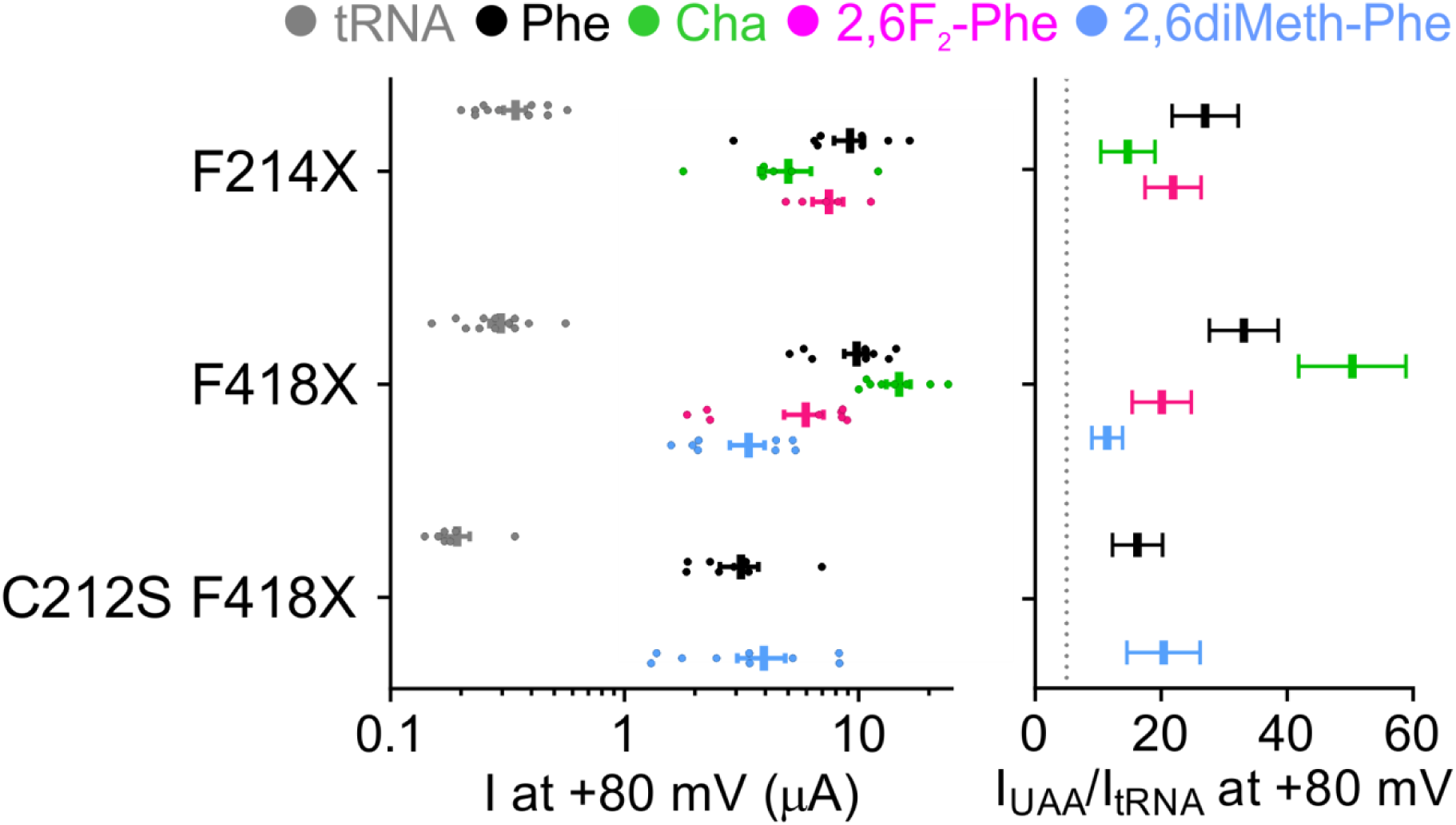
Site-specific incorporation of non-canonical amino acids into CLC-0 channels is efficient and yields robust currents. (left panel) Mean current I at +80 mV in *Xenopus laevis* oocytes for F214X, F418X and C212S F418X CLC-0 co-injected with empty tRNA (gray), Phe (black), Cha (green), 2,6F_2_-Phe (pink) and 2,6diMeth-Phe (cyan). Individual data points are shown as circles and the mean ± S.E.M. is shown as a vertical bar with error bars. (right panel). For all constructs, I_UAA_/I_tRNA_ was calculated from the means of I_UAA_ and I_tRNA_ at +80 mV as reported in the left panel. Errors were propagated. A threshold of I_UAA_/I_tRNA_>5 is imposed for specific incorporation efficiency (dashed line).

**Supplementary Table 1.**
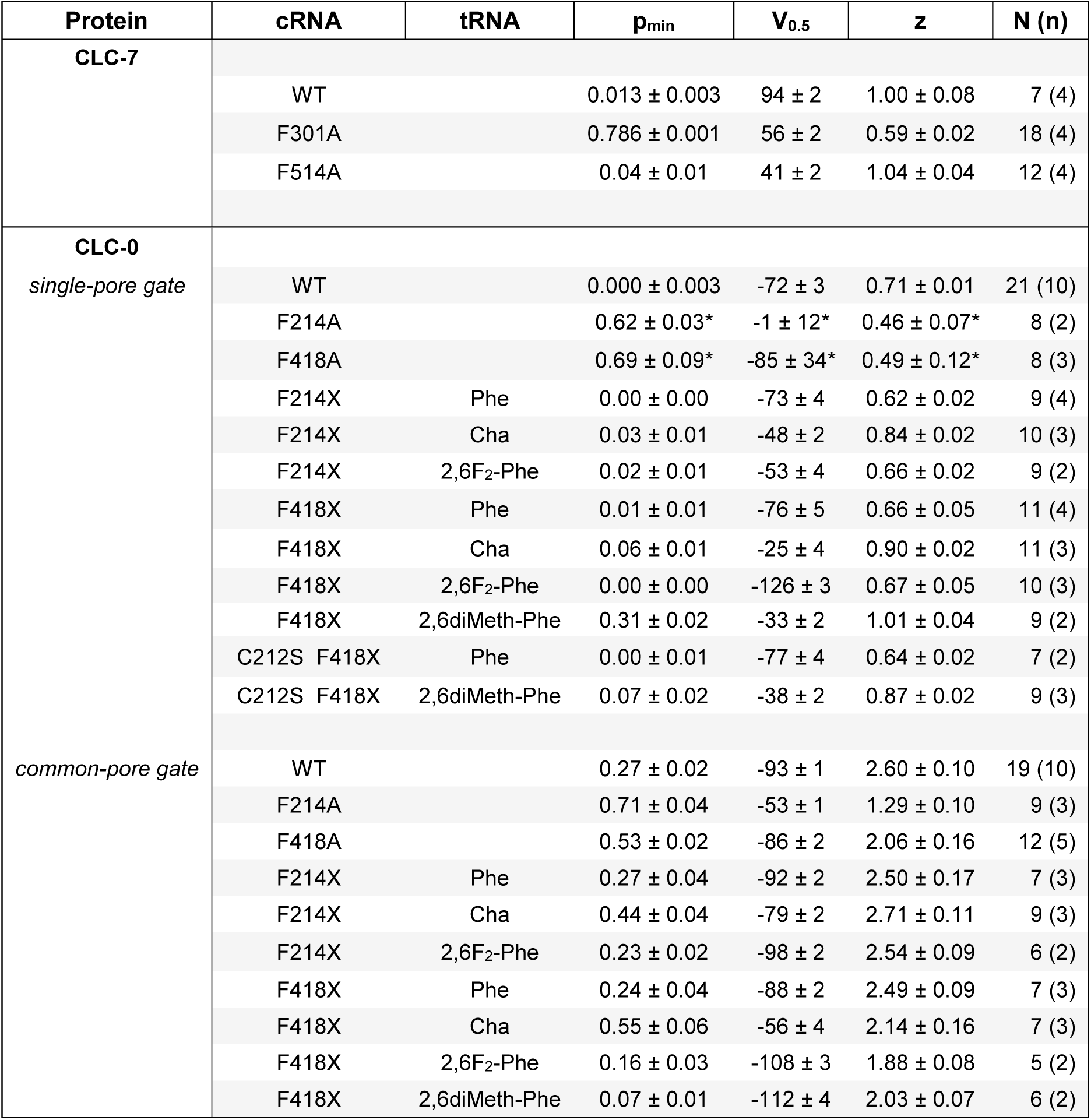
Summary of the Boltzmann fitting parameters. The following parameters from Boltzmann fits for CLC-7 and CLC-0 constructs used in Fig. 5-7 are reported as mean ± S.E.M.: *p_min_*, minimal open probability; *V_0.5_*, voltage of half maximal activation; *z*, gating charge; *N*, number of oocytes recorded; *n*, number of independent oocyte batches. * indicates values of the fit parameters that were not well constrained during fitting, as such they should be considered as estimates of the parameters.

